# A novel interplay between GEFs orchestrates Cdc42 activity during cell polarity and cytokinesis

**DOI:** 10.1101/364786

**Authors:** Brian S. Hercyk, Julie T. Rich-Robinson, Ahmad S. Mitoubsi, Marcus A. Harrell, Maitreyi E. Das

## Abstract

Cdc42, a conserved regulator of cell polarity, is activated by two GEFs, Gef1 and Scd1, in fission yeast. While *gef1* and *scd1* mutants exhibit distinct phenotypes, how they do so is unclear given that they activate the same GTPase. Using the GEF localization pattern during cytokinesis as a paradigm, we report a novel interplay between Gef1 and Scd1 that spatially modulates Cdc42. We find that Gef1 promotes Scd1 localization to the division site during cytokinesis and to the new end during polarized growth through the recruitment of the scaffold Scd2 via a Cdc42 feedforward pathway. Gef1-mediated Scd1 recruitment at the new end enables the transition from monopolar to bipolar growth. Reciprocally, Scd1 restricts Gef1 localization to prevent ectopic Cdc42 activation during cytokinesis to promote cell separation and during interphase to maintain cell shape. Our findings reveal an elegant regulatory pattern in which Gef1 establishes new sites of Scd1-mediated Cdc42 activity, while Scd1 restricts Gef1 to functional sites. We propose that crosstalk between GEFs is a conserved mechanism that orchestrates Cdc42 activation during complex cellular processes.

**Summary Statement:** Cdc42 GEFs Gef1 and Scd1 crosstalk to fine-tune Cdc42 activity. This crosstalk promotes bipolar growth and maintains cell shape in fission yeast.

## INTRODUCTION

Growth and division, fundamental processes in all cells, are essential for proper function and proliferation, and function through polarization. Cell polarization relies on the ability of the cytoskeleton to establish unique domains at the cell cortex to govern the local function and activity of specific proteins (Drubin and Nelson, 1996; Nance and Zallen, 2011). The Rho family of small GTPases serves as the primary regulators of cell polarity via actin regulation (Ridley, 2006). Active Rho GTPases bind and activate downstream targets which regulate actin cytoskeleton organization. GTPases are active when GTP-bound and inactive once they hydrolyze GTP to GDP. Guanine nucleotide Exchange Factors (GEFs) activate GTPases by promoting the binding of GTP, while GTPase Activating Proteins (GAPs) inactivate GTPases by promoting GTP hydrolysis (Bos et al., 2007). Unraveling the regulation of these GEFs and GAPs is at the crux of understanding how cell polarity is established, altered, and maintained. One conserved member of the Rho family of small GTPases, Cdc42, is a master regulator of polarized cell growth and membrane trafficking in eukaryotes (Estravis et al., 2012; Estravis et al., 2011; Etienne-Manneville, 2004; Harris and Tepass, 2010; Johnson, 1999). Like other small GTPases, Cdc42 acts as a binary molecular switch and can respond to and initiate multiple signaling pathways. In most eukaryotes, Cdc42 is regulated by numerous GEFs and GAPs, complicating our understanding of GTPase regulation (Bos et al., 2007). In the fission yeast *Schizosaccharomyces pombe*, Cdc42 is activated by two GEFs, Gef1 and Scd1 (Chang et al., 1994; Coll et al., 2003). The presence of only two Cdc42 GEFs, and the well-documented process of cell polarization in these cells, make fission yeast an excellent model system to understand the mechanistic details of cell shape establishment. Here we report that the two Cdc42 GEFs regulate each other during both cytokinesis and polarized growth. This finding provides new insights into the spatiotemporal regulation of Cdc42 during critical cellular events.

Fission yeast cells are rod-shaped and grow in a polarized manner from the two ends. These cells exhibit a unique growth pattern; cells in early G2 phase are monopolar, and grow from the old end, which existed in the previous generation. Cells transition to a bipolar growth pattern in late G2 through the process of new end take off (NETO), when growth initiates at the new end, which was formed during sister cell separation (Mitchison and Nurse, 1985). Due to this simple growth pattern, fission yeast is an excellent model to understand how a cell regulates polarized growth from multiple sites. In fission yeast, active Cdc42 displays anti-correlated oscillations between the two ends (Das et al., 2012). These oscillations arise from both positive and time-delayed negative feedback as well as from competition between the two ends (Das et al., 2012).

The regulation that gives rise to the oscillatory pattern also regulates cell dimensions and promotes bipolar growth in fission yeast. Similar Cdc42 oscillations have been observed in natural killer cells during immunological synapse formation (Carlin et al., 2011) and in budding yeast during bud emergence (Howell et al., 2012). In plant cells, the ROP GTPases show oscillatory behavior during pollen tube growth (Hwang et al., 2005). Furthermore, during migration in animal cells, the GTPases Rho, Rac, and Cdc42 are sequentially activated to enable cell protrusion (Machacek et al., 2009). These observations suggest that oscillatory behavior, which drives cell polarity, may be an intrinsic property of GTPases that is likely conserved in most organisms (Das and Verde, 2013).

Cdc42 undergoes precise spatiotemporal regulation to efficiently promote different cellular processes. In fission yeast, Cdc42 must be activated at the cell ends to promote polarized growth and restricted from the cell sides to maintain cell shape (Das et al., 2012; Das et al., 2015; Das et al., 2009). Cdc42 is also involved in cytokinesis in fission yeast. During cytokinesis, Cdc42 activation promotes septum formation, and like in other systems, Cdc42 needs to be subsequently inactivated to promote cell separation (Atkins et al., 2013; Onishi et al., 2013; Wei et al., 2016). The regulatory mechanisms that allow for these spatiotemporal activation patterns are not well understood. To explain Cdc42 activation during polarized growth, it is important to first understand how Cdc42 regulators function. Gef1 and Scd1 are partially redundant but exhibit unique phenotypes when deleted (Chang et al., 1994; Coll et al., 2003), indicating that they may regulate Cdc42 in distinct, but overlapping, manners. Scd1 oscillates between the two cell ends, much like active Cdc42 (Das et al., 2012), and is essential for polarity establishment (Chang et al., 1994). Scd1 is also required for mating and contributes to Cdc42-dependent exploration of the cell cortex (Bendezu and Martin, 2013). In contrast, *gef1* mutants are narrower and grow in a monopolar, rather than a bipolar, manner (Coll et al., 2003). Furthermore, Cdc42 activity is reduced at the new end in *gef1* mutants (Das et al., 2012). Given that Gef1 is sparsely localized to the cortex (Das et al., 2015; Tay et al., 2018) and not required for polarity establishment, it is unclear why Gef1 is required for bipolar growth. Understanding how Gef1 regulates bipolar growth will provide valuable insights into Cdc42 regulation.

Investigations into the behaviors of Gef1 and Scd1 during interphase are complicated since these GEFs overlap at sites of polarized growth. These GEFs also localize to the site of cell division during cytokinesis (Wei et al., 2016). Cytokinesis, the final step in cell division, involves the formation of an actomyosin ring that constricts, concurrent with cell wall (septum) deposition, to enable membrane ingression and furrow formation (Pollard, 2010). The temporal localization and function of the two GEFs are discernible during cytokinesis since they are recruited to the division site in succession to activate Cdc42. During cytokinesis, Gef1 localizes first to the actomyosin ring to activate Cdc42 and promote ring constriction (Wei et al., 2016). Next, Scd1 localizes to the ingressing membrane and regulates septum formation (Wei et al., 2016). The temporal difference between Gef1 and Scd1 localization at the division site allows us to investigate the significance of the GEFs in Cdc42 regulation, which is unclear from studies solely of the growing ends.

Bipolar growth occurs when the new end is able to overcome the dominance at the old end. Here we show that Gef1 enables the new end to overcome old end dominance and promote bipolar growth. We find that in *gef1* mutants both the Cdc42 GEF Scd1 and its scaffold Scd2 are localized mainly to the old ends. Using cytokinesis as a paradigm to investigate how Gef1 regulates Scd1 localization, we identify a novel crosstalk between Gef1 and Scd1. Our data indicate that Gef1 promotes the localization of Scd1 to the division site via Cdc42 activation. The scaffold Scd2 is required for Scd1 localization and binds active Cdc42 (Endo et al., 2003; Kelly and Nurse, 2011). Our data indicate that Gef1 activates Cdc42 that then recruits Scd2 and consequently Scd1. We extend these observations to the sites of polarized growth, where we show that Gef1 promotes bipolar Scd1 and Scd2 localization; indeed, Gef1 is necessary to recruit Scd1 to the non-dominant new end to initiate bipolar growth. While Gef1 promotes recruitment of Scd1 to initiate new site of Cdc42 activation, we find that Scd1 prevents ectopic Gef1 localization to the division site and the cell cortex. By this manner of regulation, Cdc42 activation is promoted at the new end of the cell with no prior growth history, but is restricted from random sites. To our knowledge, such crosstalk has not been reported to function between GEFs of the same GTPase. The interplay between the Cdc42 GEFs operates in the same manner during both cytokinesis and polarized growth, suggesting that this may be a conserved feature of Cdc42 regulation.

## RESULTS

### Gef1 enables the transition from monopolar to bipolar growth

Fission yeast transitions from monopolar to bipolar growth upon reaching a certain size (Mitchison and Nurse, 1985). It has been proposed that this size requirement is necessary to establish two stable regions of Cdc42 activity (Das et al., 2012). Additionally, since protein abundance scales with cell size, this may allow for the accumulation of sufficient GEFs and other polarity factors to maintain two sites of Cdc42 activity (Das et al., 2012). As per these models, it has been suggested that in *gef1Δ* cells the total active Cdc42 levels are insufficient to allow bipolar growth, thus resulting in monopolarity. However, the dependence of bipolarity on cell size and protein abundance cannot be explained in G1-arrested *cdc10-129* mutants (Marks et al., 1986). *cdc10-129* cells shifted to 36°C remain monopolar even as they grow longer compared to *cdc10+* cells (Figure 1A). Alternatively, it is possible that bipolar growth requires intricate regulation of Cdc42 activation at the new end, and that Gef1 is specifically involved in this process.

**Figure 1:**
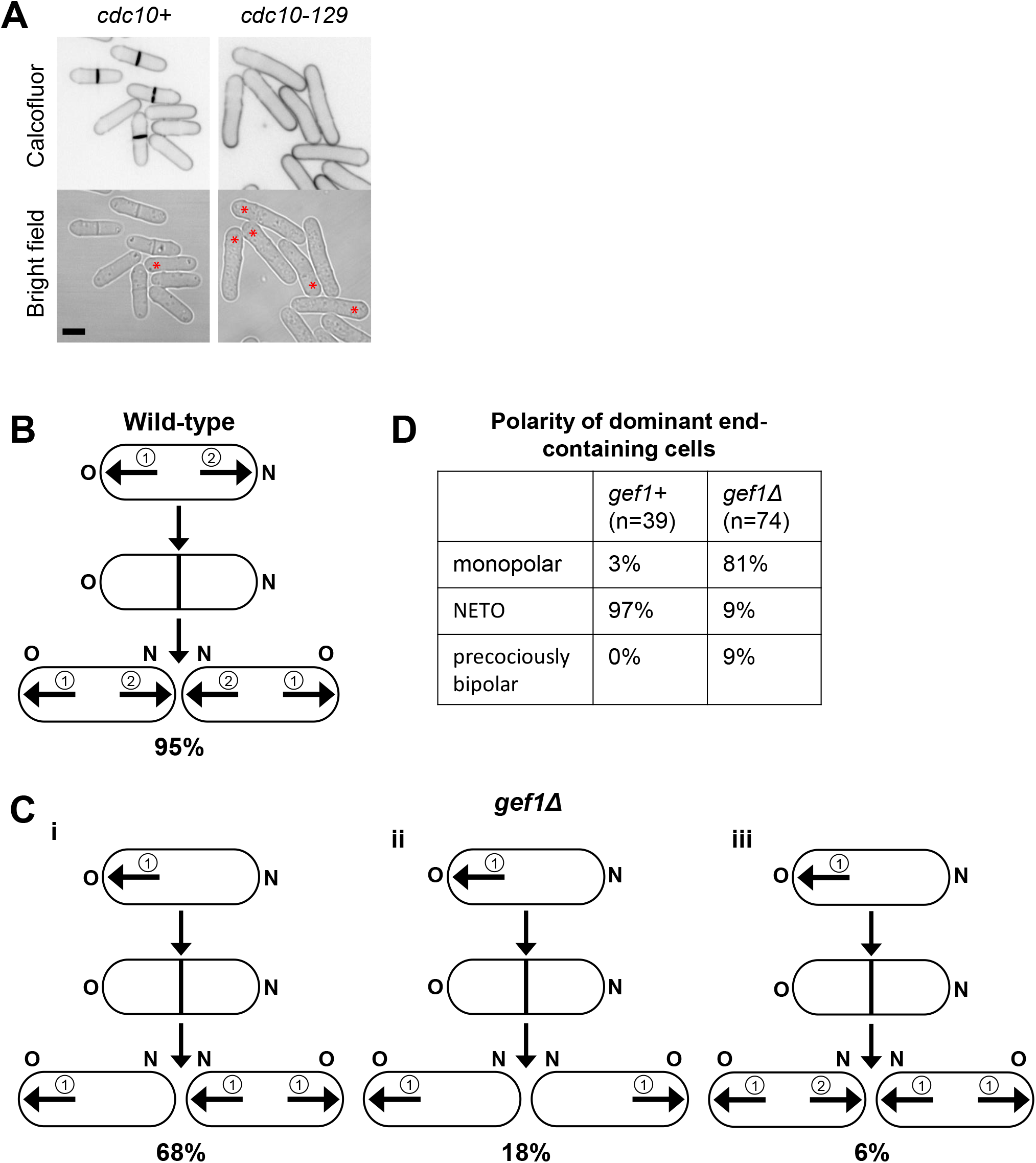
Gef1 promotes bipolar growth via new-end-take-off. **(A)** Polarized growth phenotypes of calcofluor stained *cdc10+* and *cdc10-129* cells grown at 36°C. *cdc10+* cells initiate bipolar growth, while *cdc10-129* cells arrest at G1-phase and remain monopolar. Red asterisks mark cells that exhibit monopolar growth. **(B)** Wild-type cells predominately display old end growth followed by a delayed onset of new-end growth. **(C)** i. In *gef1Δ*, 68% of monopolar cells yield a monopolar daughter cell from the end that grew in the previous generation and a bipolar daughter cell from the end that failed to grow in the previous generation. ii.18% of monopolar cells yield two monopolar cells. iii. 6% of monopolar cells yield two monopolar cells. Circled numbers describe the order of growth. Arrows correspond to direction of growth. **(D)** Quantification of cell growth pattern in *gef1+* and *gef1Δ* cells. NETO-new end take off, Scale bar = 5μm.

We examined the growth patterns of *gef1Δ* mutants to gain insight into the transition to bipolar growth. The old ends in fission yeast initiate growth immediately after completion of division and cell separation. As the cell elongates, it eventually begins to grow at the new end, resulting in bipolar growth (Figure 1B) (Mitchison and Nurse, 1985). The two ends in fission yeast compete for active Cdc42; at first the old end wins this competition (Das et al., 2012). The old end can thus be said to be dominant over the new end in a newborn cell, and always initiates growth first. The new end must overcome the old end’s dominance in order to initiate its growth.

We find that 68% of monopolar *gef1Δ* mutant cells exhibit a growth pattern in which one daughter cell is monopolar and the other daughter cell is prematurely bipolar (Figures 1C and S1). In monopolar *gef1Δ* cells, growth predominantly occurs at the old end, which grew in the previous generation (Figures 1C and S1). In these monopolar cells, the new end frequently fails to grow since it cannot overcome the old end’s dominance. The daughter cell that inherits its parent cell’s non-growing end typically displays precocious bipolar growth, indicating that these cells do not contain a dominant end. Our data suggest that for a cell end to be dominant it needs to have grown in the previous generation. These results indicate that the new ends of *gef1Δ* cells are not well-equipped to overcome old end dominance. Indeed, we find that in *gef1+* cells, 97% of daughter cells derived from a growing end display a normal growth pattern in which new end take-off (NETO) occurs only after the old end initiates growth (Figure 1C,D). In *gef1Δ* cells, only 9% of daughter cells derived from a growing end display NETO; instead, 81% of daughter cells derived from a growing end failed to initiate growth at their new end and were thus monopolar (Figure 1C,D). These data reveal that Gef1 enables the new end to overcome old end dominance to promote bipolar growth.

### Gef1 enables bipolar localization of Scd1 and Scd2 to the cell poles

To address the mechanism through which Gef1 enables bipolar growth, we examined other polarity factors that promote Cdc42 activation, Scd1 and Scd2. Scd1 and Scd2, like active Cdc42, undergo oscillations between the two competing ends (Das et al., 2012); thus, a cell undergoing bipolar growth does not always display bipolar Scd1 or Scd2 localization. We find that fewer new ends in *gef1Δ* cells exhibited Scd1-3xGFP; bipolar Scd1-3xGFP was observed in 30% of interphase *gef1+* cells, but only in 14% of *gef1Δ* cells (Figure 2A,B; p=0.0004). Similarly, we observed a decrease in bipolar Scd2 in cells lacking *gef1;* 70% of *gef1+* cells displayed bipolar Scd2-GFP localization, but this was reduced to 30% in *gef1Δ* cells (Figure 2A,B, p<0.0001). While these data suggest that Gef1 promotes Scd1 and Scd2 localization to the new end to enable bipolar growth, this interpretation suffers from some complications. The old and new ends compete with each other for active Cdc42 (Das et al., 2012). It is possible that in *gef1Δ* mutants the new end fails to grow since the old end traps Scd1 and Scd2, resulting in monopolar distribution of these proteins. This would suggest that Gef1 removes Scd1 and Scd2 from the old end, thereby making it available for the new end. However, we did not find enhanced Scd1-3xGFP levels at the old end in *gef1Δ* cells compared to *gef1+* cells (Figure S2). This suggests that Gef1 does not prevent accumulation of Scd1 and Scd2 at the old ends. Alternately, it is possible that Gef1 and Scd1 act independently to activate Cdc42 that the old and new ends compete for. As the level of total active Cdc42 increases, it saturates the dominant old end and the new end is now able to initiate growth and localize Scd1 and Scd2. In this scenario, Scd1 and Scd2 localization at the new end would be an indirect effect of increased Cdc42 activity and loss of competition at the old end. To distinguish whether Scd1 and Scd2 localization at the new end requires Gef1, or whether it is an indirect outcome simply due to saturation of active Cdc42 at the old end, we turned to the division site. The division site does not compete with any other site in the cell for active Cdc42. Moreover, the localization of the GEFs can be temporally resolved at this site allowing us to examine the relationship between Gef1, Scd1, and Scd2.

**Figure 2:**
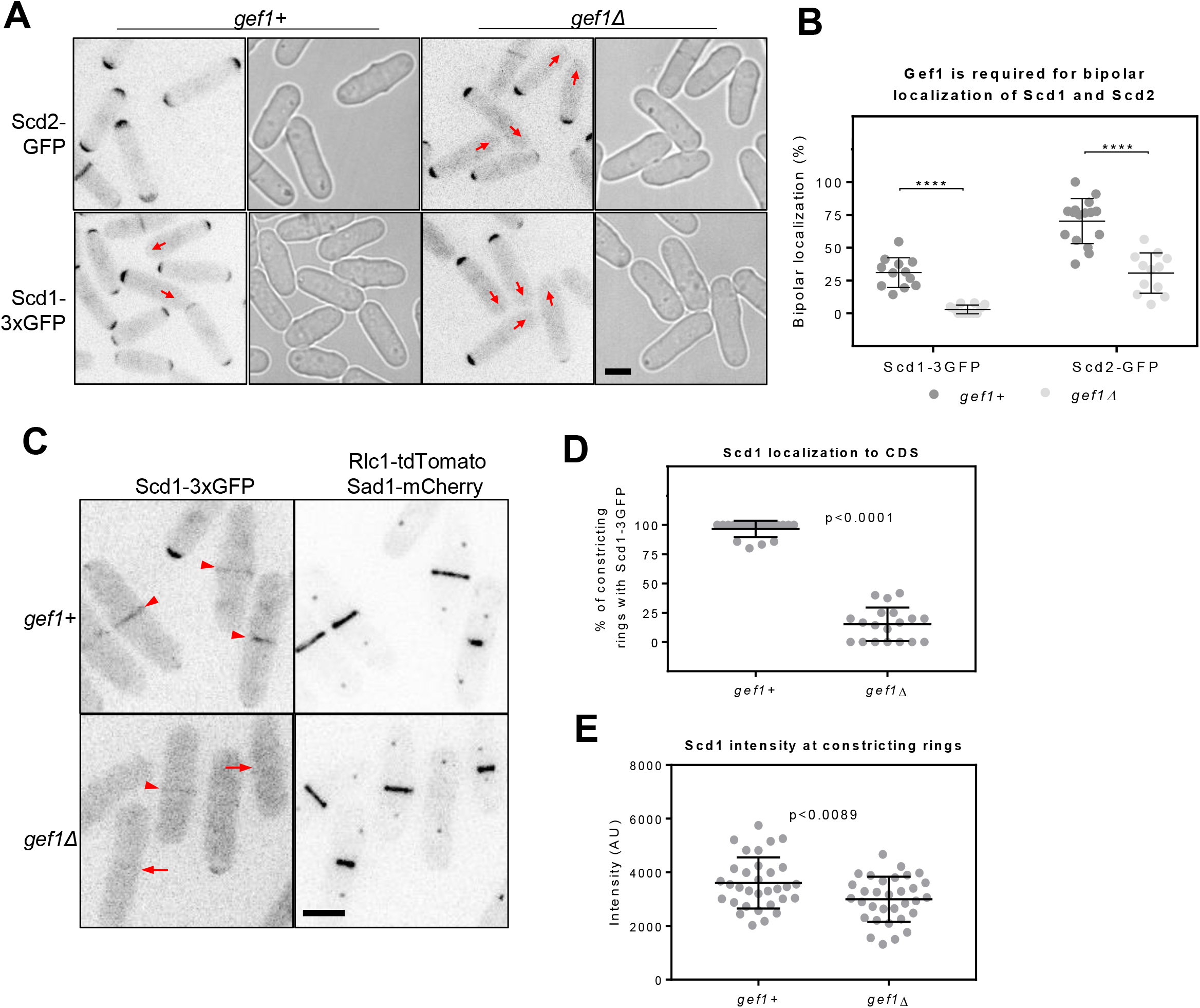
Gef1 promotes localization of polarity factors Scd1 and Scd2 to the new end. **(A)** Scd2-GFP and Scd1-3xGFP localization to the sites of polarized growth in *gef1+* and *gef1Δ* cells. Red arrows indicate the new ends of monopolar cells that do not recruit Scd2-GFP or Scd1-3xGFP. **(B)** Quantifications of bipolar Scd1-3xGFP and Scd2-GFP localization in the indicated genotypes. Significance determined by one-way ANOVA with Tukey’s multiple comparisons post hoc test (****, p<0.0001). **(C)** Scd1-3xGFP localization in *gef1+* and *gef1Δ* cells expressing the ring and SPB markers Rlc1-tdTomato and Sad1-mCherry respectively. Arrowheads label cells with Scd1-3xGFP localized to the division site, while arrows mark cells with constricting rings that lack Scd1-3xGFP at the division site. **(D and E)** Quantification of Scd1-3xGFP localization to and intensity at constricting rings in the indicated genotypes. Reported p values from Student’s t-tests. All data points are plotted in each graph, with black bars on top of data points that show the mean and standard deviation for each genotype. All images are inverted max projections with the exception of bright field. CDS, Cell division site. Scale bar = 5μm.

### Gef1 promotes Scd1 recruitment to the division site

We have reported that Scd1 localizes to the membrane adjacent to the actomyosin ring after Gef1 to activate Cdc42 along the membrane barrier (Wei et al., 2016). Since Scd1 arrives at the division site soon after Gef1, it is possible that Gef1 promotes Scd1 localization. Alternately, if Gef1 and Scd1 act independently, then loss of *gef1* would not impact Scd1 localization to the division site. To test this, we examined whether Scd1 localization to division site is Gef1-dependent. Both Gef1 and Scd1 are low-abundance proteins and are not suitable for live cell imaging over time. This complicates the investigation of the temporal localization of these proteins. To overcome this limitation, we used the actomyosin ring as a temporal marker. The actomyosin ring undergoes visibly distinct phases during cytokinesis: assembly, maturation, constriction, and disassembly. We determined the timing of protein localization to the division site by comparing it to the corresponding phase of the actomyosin ring. We have previously reported that ring constriction is delayed in *gef1Δ* mutants (Wei et al., 2016). To eliminate any bias in protein localization due to this delay, we only analyzed cells in which the rings had initiated constriction. In *gef1Δ* mutants, the number of constricting rings that recruited Scd1-3xGFP decreased to 15% from 96% in *gef1+* (Figure 2C,D, p<0.0001). Furthermore, the *gef1Δ* cells that managed to recruit Scd1-3xGFP did not do so as efficiently as *gef1+* cells, given the 15% decrease in Scd1-3xGFP fluorescence intensity at the division site (Figure 2C,E, p=0.0089). This indicates that Gef1 promotes Scd1 localization to the division site and that the two GEFs are not independent.

### Cdc42G12V is sufficient to restore Scd1 to the CDS in *gef1Δ*

Next, we investigated how Gef1 promotes Scd1 recruitment to the division site. GEF recruitment to sites of Cdc42 activity occurs via positive feedback, as reported in budding yeast (Butty et al., 2002; Irazoqui et al., 2003; Kozubowski et al., 2008). In this model, activation of Cdc42 leads to further recruitment of the scaffold BEM1, which then recruits the GEF CDC24 to the site of activity, thus helping to break symmetry and promote polarized growth. A similar positive feedback may also exist in fission yeast (Das et al., 2012; Das and Verde, 2013). We hypothesized that Gef1-activated Cdc42 acts as a seed for Scd1 recruitment to the division site. To test this, we asked whether constitutive activation of Cdc42 could rescue the Scd1 recruitment defect exhibited by *gef1Δ.* In order for this approach to work, the constitutively active Cdc42 must localize to the division site. Localization of active Cdc42 is visualized via the bio-probe CRIB-3xGFP that specifically binds GTP-Cdc42. Since our previous work reported that Cdc42 activity is reduced at the division site in *gef1Δ* cells (Wei et al., 2016), we validated this approach by first testing whether constitutively active Cdc42 restores CRIB-3xGFP localization at the division site in *gef1Δ* cells. The constitutively active allele *cdc42G12V* and the bio-probe CRIB-3xGFP were expressed in *gef1+* and *gef1Δ* cells. Mild expression of *cdc42G12V* was sufficient to restore CRIB-3xGFP intensity at the division site to physiological levels in *gef1Δ* (Figure 3A,B, p<0.0001). Likewise, expression of *cdc42G12V* restored Scd1-tdTomato localization to the division site in *cdc42G12V gef1Δ* cells (Figure 3C,D). This demonstrates that active Cdc42 alone is sufficient to recruit Scd1. Next, we asked how does active Cdc42 promote Scd1 localization. We examined downstream targets of active Cdc42 for this purpose. The Cdc42 ternary complex consists of the GEF Scd1, the scaffold protein Scd2, and the downstream effector Pak1 kinase (Endo et al., 2003). Observations in budding yeast suggest that the PAK kinase may mediate GEF recruitment (Kozubowski et al., 2008). Contrary to this hypothesis, we find that Scd1-3xGFP intensity increases in the *nmt1 switch-off* mutant allele of *pak1*, compared to *pak1+* cells (Figure S3). These findings support similar observations reported in the hypomorphic temperature-sensitive *pak1* allele, *orb2-34* (Das et al., 2012), and indicate that *pak1* does not facilitate Scd1 recruitment to the site of action.

**Figure 3:**
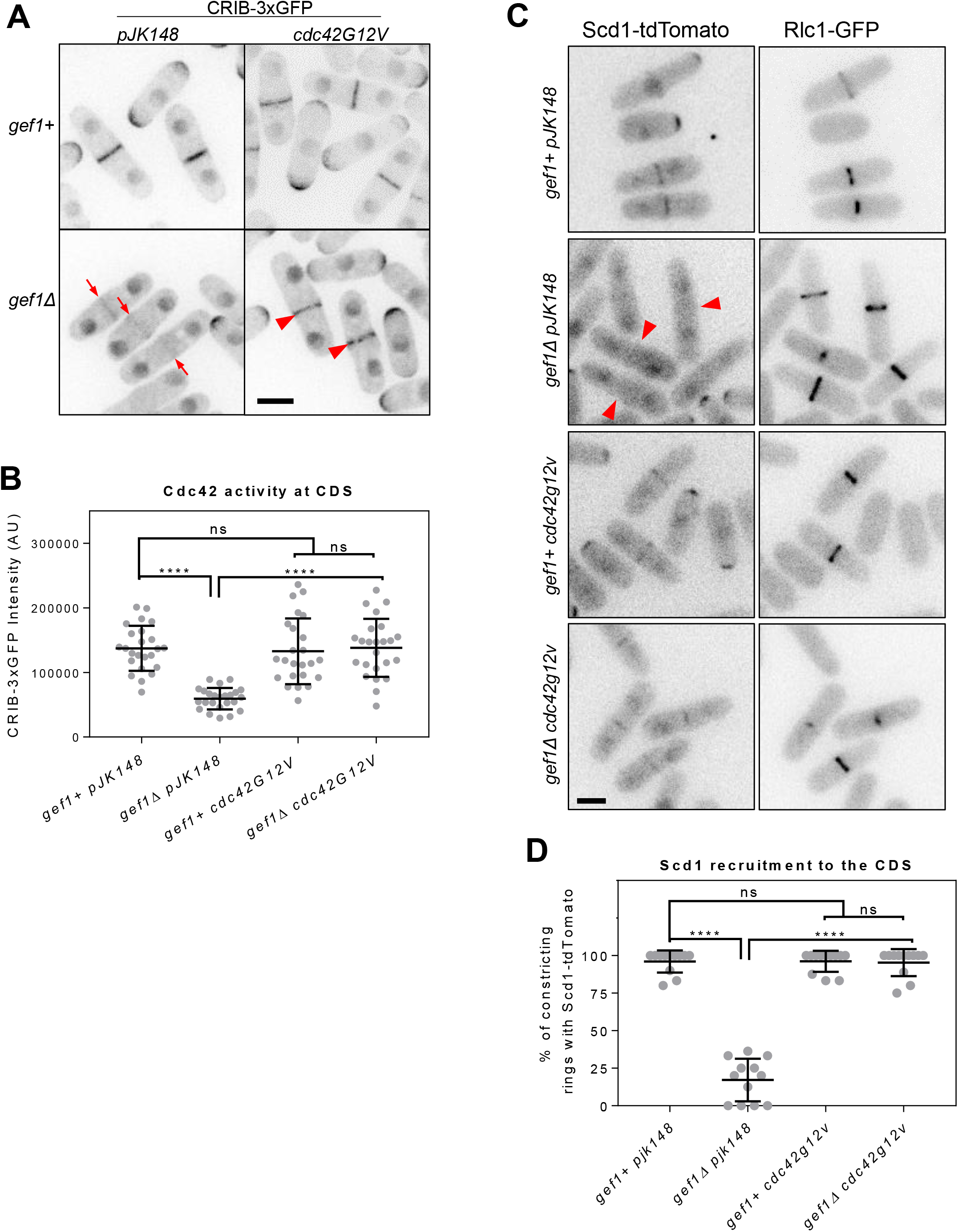
Gef1-mediated Cdc42 activation recruits Scd1 to the division site. **(A)** CRIB-3xGFP localization to the division site in *gef1+* and *gef1Δ* cells transformed with the control vector *pJK148* or *cdc42G12V.* Red arrows indicate cells with reduced Cdc42 activity at the division site in control *gef1Δ* cells, while red arrowheads indicate restored Cdc42 activity at the division site in *gef1Δ* cells expressing cdc42G12V. **(B)** Quantification of CRIB-3xGFP intensity at the cell division site (CDS) in the indicated genotypes (****, p<0.0001). **(C)** Scd1-tdTomato localization to constricting rings marked by Rlc1-GFP in *gef1+* and *gef1Δ* cells transformed with the control vector *pJK148* or *cdc42G12V.* Red arrows indicate cells with constricting rings lacking Scd1-tdTomato in control *gef1Δ* cells, while red arrowheads indicate restored Scd1-tdTomato at the constricting rings of *gef1Δ* cells expressing cdc42G12V. **(D)** Quantification of cells with Scd1-tdTomato at the constricting rings of cells of the indicated genotypes (****, p<0.0001). All data points are plotted in each graph, with black bars on top of data points that show the mean and standard deviation for each genotype. All images are inverted max projections. Significance determined by one-way ANOVA with Tukey’s multiple comparisons post hoc test. Scale bars = 5μm.

### Gef1 promotes Scd2 localization to the division site, which in turn recruits Scd1

Scd2 is a component of the Cdc42 ternary complex, and binds active Cdc42 (Endo et al., 2003; Wheatley and Rittinger, 2005). We hypothesized that Gef1-dependent active Cdc42 recruits Scd1 to the division site through the scaffold Scd2. If this were true, Scd2-GFP localization to the division site should be Gef1-dependent. Indeed, we find that *gef1Δ* cells displayed a significant decrease in Scd2-GFP-containing assembled rings compared to *gef1+* cells. In *gef1Δ* mutants, the number of rings that recruited Scd2-GFP prior to ring constriction decreased to 8% compared to 88% in *gef1+*, indicating a delay in Scd2 recruitment (Figure 4A,B, p>0.0001). Although *gef1Δ* cells were able to recruit Scd2 to the division site once ring constriction began, the fluorescence intensity of Scd2-GFP at the division site was reduced by 61% compared to *gef1+* cells (Figure 4A,C, p>0.0001, Figure S4). Gef1 thus promotes Scd2 localization to the division site. Since previous work indicates that Scd1 and Scd2 require each other for their localization (Kelly and Nurse, 2011), it is possible that a decrease in Scd2 at the division site observed in *gef1* mutants is due to a decrease in Scd1 at this site. However, contrary to previous findings, we observed that Scd2-GFP localization at the division site is not impaired in *scd1Δ* cells (Figure 4D). In contrast, Scd1-3xGFP localization is completely abolished at the division site in *scd2Δ* cells (Figure 4E). We find that Scd1 requires Scd2 for its localization to the division site. Scd2 localization, on the other hand, is independent of Scd1 and depends on Gef1 instead.

**Figure 4:**
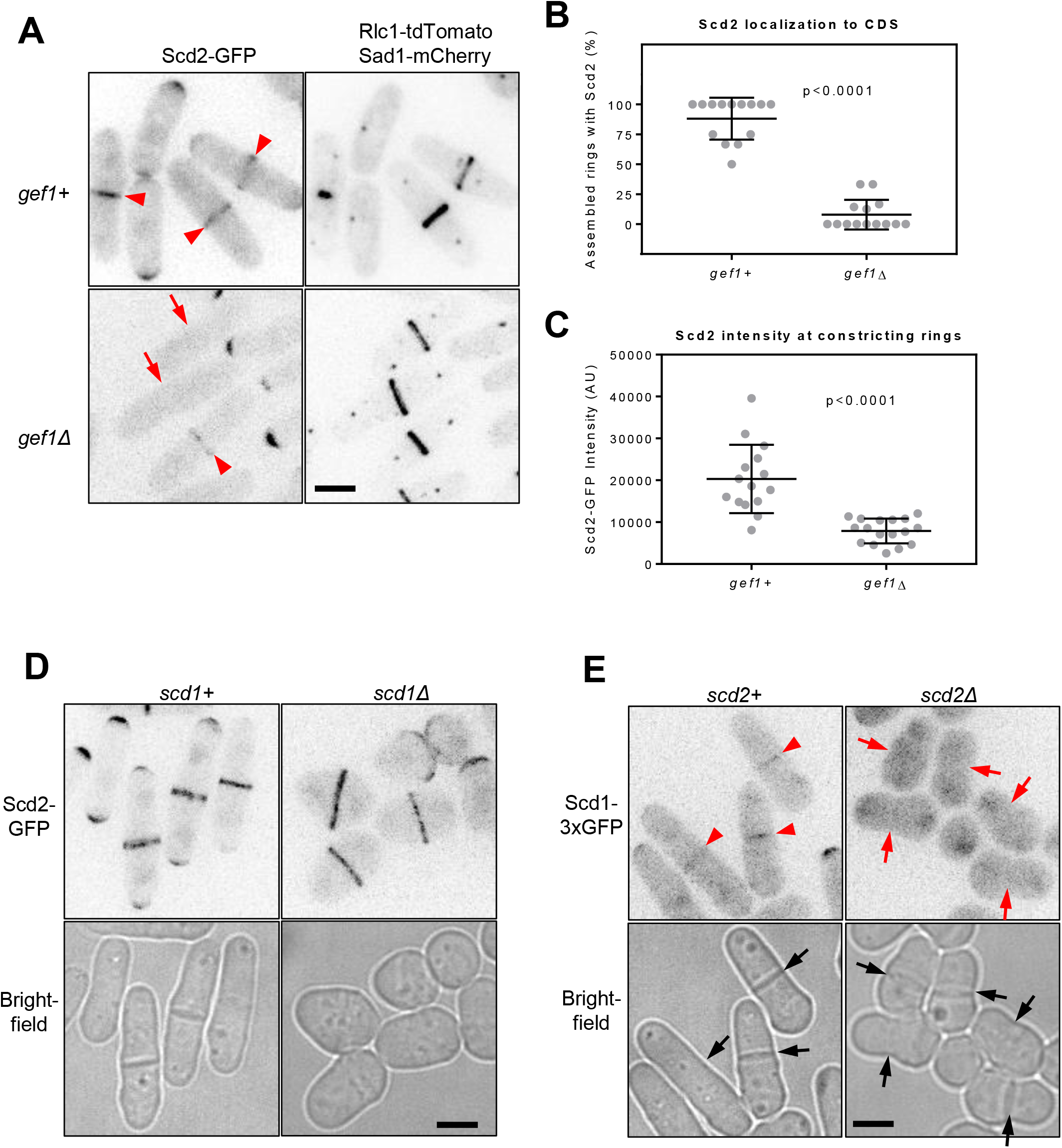
Gef1 promotes Scd1 and Scd2 localization to the division site. **(A)** Scd2-GFP localization in *gef1+* and *gef1Δ* cells expressing the ring and SPB markers Rlc1-tdTomato and Sad1-mCherry. Arrowheads label cells with Scd2-GFP localized to constricting rings, while arrows mark cells with assembled rings that lack Scd2-GFP at the division site. **(B and C)** Quantification of Scd2-GFP localization and intensity in the indicated genotypes. All data points are plotted in each graph, with black bars on top of data points that show the mean and standard deviation for each genotype. All images are inverted max projections with the exception of bright field. Significance determined by student’s t-test. **(D)** Scd2-GFP localization to the division site in *scd1+* and *scd1Δ* cells. **(E)** Scd1-3xGFP localization in *scd2+* and *scd2Δ* cells. Division site marked by black arrows in the bright field images. Scd1-3xGFP localization to the division site indicated by red arrowheads. Red arrows show absence of Scd1-3xGFP at the division site. All images are inverted max projections with the exception of bright field. Scale bar = 5μm.

Altogether, our data reveal that Gef1 promotes Scd2 localization to the division site, which then recruits Scd1. Based on these data, we hypothesized that Gef1, Scd2, and Scd1 sequentially localize to the division site. To test this, we examined the temporal localization of these proteins to the division site. Since these proteins do not lend themselves to extended time lapse imaging, we used the spindle pole bodies as an internal timer. The distance between spindle pole bodies is a well-established temporal marker to determine a cell’s cytokinetic stage. The spindle pole body distance increases as mitosis progresses until the cell reaches anaphase B (Nabeshima et al., 1998), at which time the actomyosin ring starts to constrict (Wu et al., 2003). The distance between the two spindle pole bodies is thus used as an internal clock that helps to time the recruitment of other proteins (Nabeshima et al., 1998). We acquired numerous still images and calculated the distance between the spindle pole bodies, marked by Sad1-mCherry, during anaphase A or anaphase B. We report the spindle pole body distance at which Gef1-mNG (monomeric NeonGreen), Scd1-3xGFP, and Scd2-GFP signals are visible at the division site before the actomyosin rings starts constriction (Figure 5A). Next, we calculated the spindle pole body distance at which Gef1-mNG, or Scd2-GFP or Scd1-3xGFP were observed at the division site. We find that Gef1-mNG and Scd2-GFP appear around the same time during mitosis with a mean spindle pole body distance of 5.9μm, and 6.6 μm, respectively (Figure 5B. not significant). Scd1-3xGFP arrives later with a longer mean spindle pole body distance of 7.6μm (Figure 5B. p=0.005). Next, we employed dual color imaging of cells expressing two fluorescently tagged proteins to validate the findings above. In cells expressing both Gef1-tdTomato and Scd2-GFP, 100% of assembled rings with Gef1 invariably also had Scd2 at the division site (Figure 5C,D). In cells expressing Scd2-GFP and Scd1-tdTomato, 59% of assembling rings localized both Scd2 and Scd1, while 41% contained only Scd2 (Figure 5C,D). In cells expressing both Gef1-tdTomato and Scd1-3xGFP, 61% of assembling rings with Gef1 also had Scd1, while 39% contained only Gef1 (Figure 5C,D). Altogether, this demonstrates that Gef1 recruits Scd2 to ring prior to Scd1.

**Figure 5:**
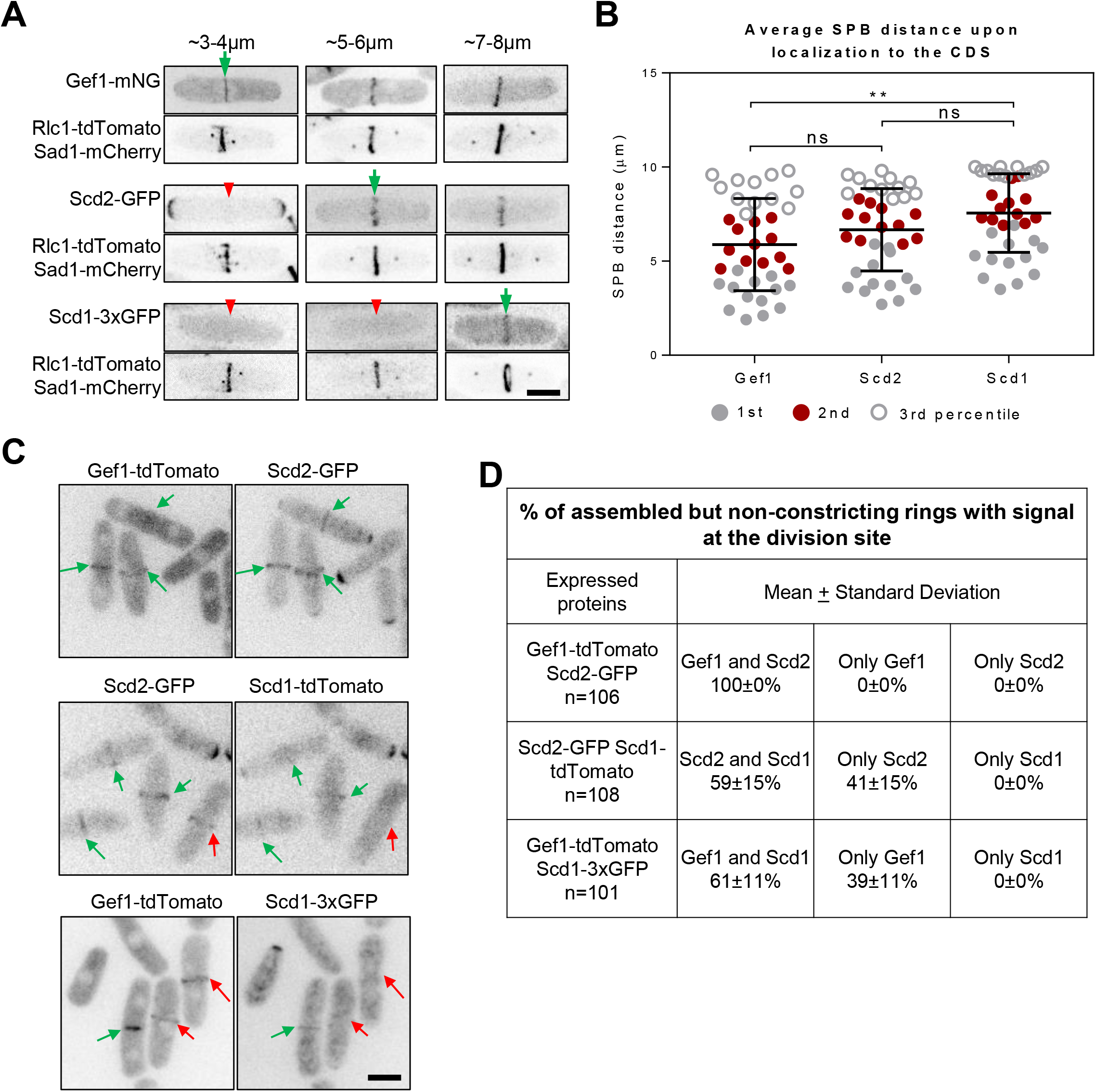
Temporal localization of Gef1, Scd2, and Scd1 to the site of cell division. **(A)** Representative images showing the localizations of Gef1-mNG, Scd2-GFP, and Scd1-3xGFP (top panels) as a function of spindle pole body distance (bottom panels). The range of SPB distance is listed for each column. Green arrows indicate the earliest time point at which signal is visible. Red arrowheads indicate time points prior to localization. **(B)** Quantification of Gef1, Scd2, and Scd1 localization to the division site in a temporal manner, showing the means of the distance between spindle poles of the first 33^rd^ percentile of early anaphase cells at which signal first appears (grey circles, Significance determined by one-way ANOVA with Tukey’s multiple comparisons post hoc test. **, p=0.005). Second and third percentiles shown in red circles and grey hollows, respectively. **(C)** Representative images of early anaphase cells expressing combinations of Scd1-3xGFP, Gef1-tdTomato, Scd2-GFP, and Scd1-tdTomato. Co-localization is marked by green arrow, and red arrows mark sites with only one kind of protein **(D)** Table summarizing the order in which Gef1, Scd2, and Scd1 localize to the division site. All data points are plotted in each graph, with black bars on top of data points that show the mean and standard deviation for each genotype. All images are inverted max projections. Scale bars = 5μm. CDS, Cell Division Site.

### Gef1 recruits Scd1 to the new end via Cdc42 activation

Next, we asked if Gef1 recruits Scd1 to the new end via Cdc42 activation, just as it does at the division site. We first tested whether expression of constitutively active Cdc42 results in bipolar localization of active Cdc42 in *gef1Δ* cells, as indicated by CRIB-3xGFP localization. Low-level expression of *cdc42G12V* was sufficient to restore bipolar CRIB-3xGFP localization in *gef1Δ*, compared to the empty-vector-containing *gef1Δ* mutants (Figure 6A,B; p<0.0001). We observed bipolar CRIB-3xGFP in 35% of *gef1+* cells transformed with the empty vector and in 36% of cells expressing *cdc42G12V.* In *gef1Δ* mutants transformed with the empty vector, we observed bipolar CRIB-3xGFP in only 8% of cells. In contrast, in *gef1Δ* mutants, low levels of *cdc42G12V* expression restored bipolar CRIB-3xGFP to 34%. In the same cells, we observed bipolar Scd1-tdTomato in 21% of *gef1+* cells transformed with the empty vector, and in 23% of cells expressing *cdc42G12V.* In *gef1Δ* mutants transformed with the empty vector, we observed bipolar Scd1-tdTomato in only 6% of cells. Further, in *gef1Δ* mutants expressing low levels of *cdc42G12V*, bipolar Scd1-tdTomato was restored to 19% (Figure 6A,C). Thus, expression of *cdc42G12V* was sufficient to restore bipolar Scd1-tdTomato localization to the cell ends in *gef1Δ* mutants, just as it was at the division site (Figure 6A,C). This demonstrates that Gef1 promotes Scd1 localization to the new ends through Cdc42 activation. As a further test, we asked whether Scd1 localization would be enhanced at the new end in the precociously bipolar *gef1S112A* mutant, in which Gef1S112A constitutively localizes to the cortex. Indeed, we find that 52% of *gef1S112A* cells show bipolar Scd1-3xGFP localization, while this is seen in only 33% of *gef1+* cells (Figure S5; p<0.0001). At the division site, Gef1 colocalizes with the scaffold Scd2, which then recruits Scd1. While Scd1 requires Scd2 for its localization, Scd2 localization is independent of Scd1 (Figure S6). Scd2 binds active Cdc42, so it is possible that Cdc42 activation mediated by Gef1 leads to Scd2 recruitment. To test this, we monitored ectopic Gef1 localization in fission yeast cells. In cells treated with the actin-depolymerizing drug Latrunculin A (LatA) Gef1-tdTomato ectopically localizes to the cell sides (Figure 6D). In these cells, Scd2-GFP colocalizes with ectopic Gef1-tdTomato at the cell sides. Moreover, in *gef1Δ* cells Scd2-GFP fails to localize to the cell sides upon LatA treatment (Figure 6E). In these cells, Scd2-GFP remains at the cell ends, but the levels are much reduced. This provides further evidence that Gef1 marks the site for Scd2 recruitment at the cell cortex.

**Figure 6:**
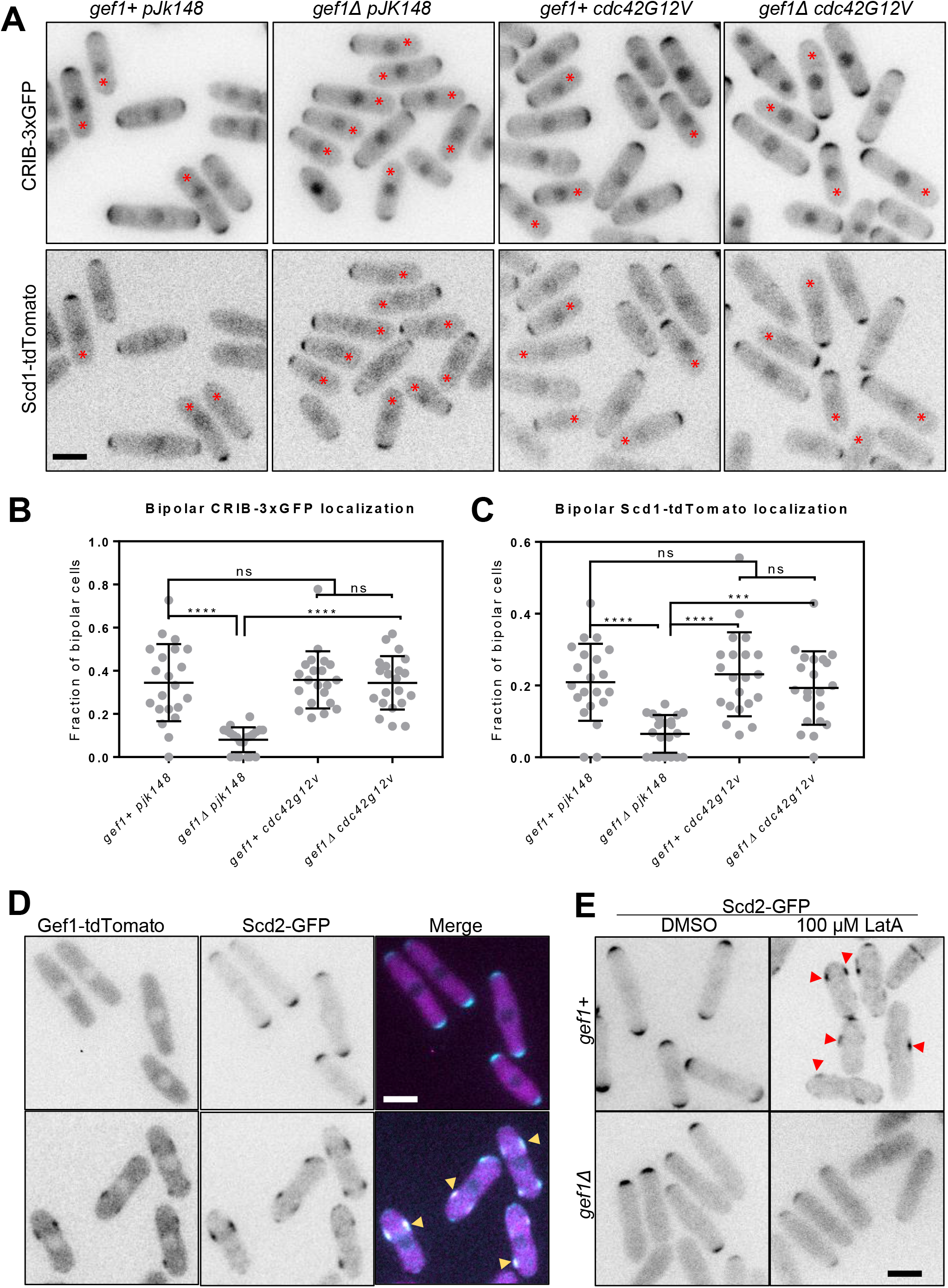
Gef1 promotes Scd1 localization to the new end. **(A)** CRIB-3xGFP and Scd1-tdTomato localization at cell tips, and restoration of bipolar growth in *gef1+* and *gef1Δ* cells transformed with the control vector *pJK148* or *cdc42G12V.* Red asterisk indicate the new end of monopolar cells. **(B and C)** Quantification of the percent of cells that exhibit bipolar CRIB-3xGFP and Scd1-tdTomato localization at cell tips in the indicated genotypes (Significance determined by one-way ANOVA with Tukey’s multiple comparisons post hoc test. ***, p<0.001, ****, p<0.0001). All data points are plotted in each graph, with black bars on top of data points that show the mean and standard deviation for each genotype. All images are inverted max projections. **(D)** Localization of Gef1-tdTomato and Scd2-GFPin DMSO or LatA treated cells. **(E)** Localization of Scd2-GFP in *gef1+* and *gef1Δ* cells treated with DMSO or LatA. All images are inverted max projections with the exception of bright field unless specified. Scale bar = 5μm.

### Scd1 prevents ectopic Gef1 localization to the cell cortex and the division site

Cells treated with LatA show ectopic Cdc42 activation at the cell sides (Mutavchiev et al., 2016). We find that LatA treatment leads to ectopic Gef1 and Scd2 localization to the cell sides. As reported earlier, we find that LatA treatment leads to a severe loss in Scd1-mNG levels at the cell ends and at the site of cell division (Figure S7). We asked if ectopic Gef1 localization in LatA-treated cells is due to loss of Scd1 from the cell cortex. To test this, we analyzed Gef1-mNG localization in *scd1Δ* mutants. We find that in *scd1+* cells, Gef1-mNG is mostly cytoplasmic and displays sparse but polarized localization at cell ends (Figure 7Ai). In *scd1Δ* mutants, Gef1-mNG shows depolarized cortical localization with random patches all over the cortex similar to that in *scd1+* cells treated with LatA (Figure 7Aii,v). Gef1-mNG localization in *scd1Δ* cells treated with LatA is similar to that in untreated *scd1Δ* cells. This suggests that Scd1 is required to prevent ectopic Gef1 localization and restricts it to the cell ends.

**Figure 7:**
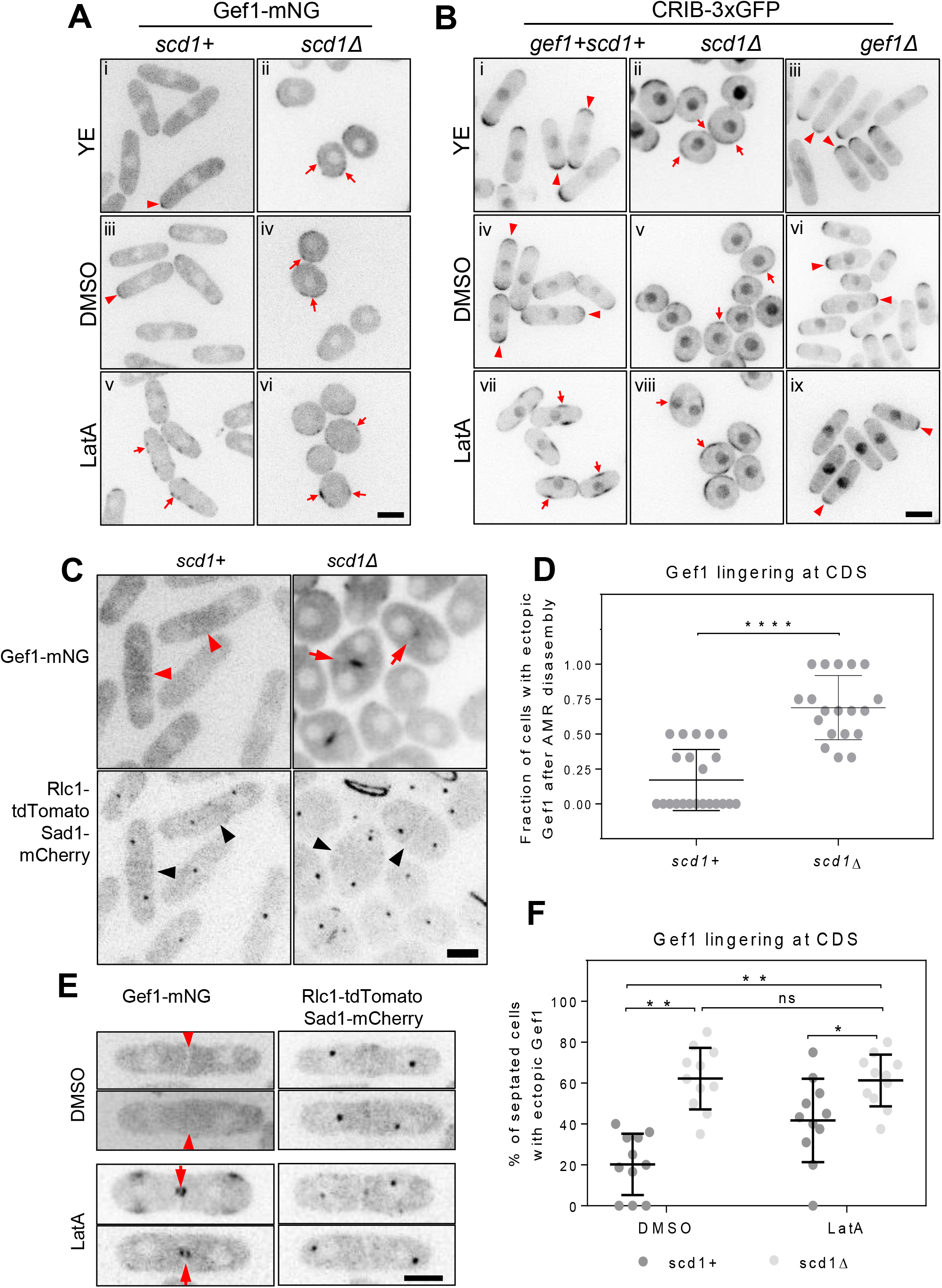
Scd1 and actin prevent ectopic Gef1 localization to promote polarized growth, and promote Gef1 removal from the division site after ring constriction. **(A)** Gef1-mNG localization in *scd1+* and *scd1Δ* cells grown in yeast extract (YE, top panel) or treated with DMSO (middle panel) or 10μM LatA (bottom panel). **(B)** CRIB-3xGFP localization in *gef1+* and *gef1Δ* cells treated with DMSO (top panel) or 10μM LatA (bottom panel). Arrowheads indicate cells with CRIB-3xGFP or Gef1-mNG localized to regions of polarized growth. Arrows indicate cells with CRIB-3xGFP or Gef1-mNG localized to non-polarized regions on the cell cortex. Images in (A) and (B) are inverted max projections of the medial 4-7 cell slices. **(C)** Gef1-mNG localization to the division site after ring disassembly in *scd1+* and *scd1Δ* cells expressing the ring and SPB markers Rlc1-tdTomato and Sad1-mCherry. Black arrowheads mark the membrane barrier in cells post-ring disassembly. Red arrowheads mark cells post-ring disassembly that lack Gef1-mNG at the membrane barrier. Red arrows mark cells with Gef1-mNG localized to the membrane barrier post-ring assembly. **(D)** Quantification of Gef1 lingering at the division site in *scd1+* and *scd1Δ* cells (****, p<0.0001, student’s t-test). **(E)** Gef1-mNG localization in septated cells expressing the ring and SPB markers Rlc1-tdTomato and Sad1-mCherry, treated with either DMSO, 10μM LatA, or 100μM CK666. Red arrowheads mark cells post-ring disassembly that lack Gef1-mNG at the membrane barrier. Red arrows indicate cells with Gef1-mNG localized to the membrane barrier post-ring assembly. Red arrowheads indicate cells with absence of Gef1-mNG to the membrane barrier. **(F)** Quantification of Gef1 lingering at the division site in septated *scd1+* and *scd1Δ* cells treated with 10μM LatA or DMSO (*, p<0.05. **,p<0.01, one-way ANOVA with Tukey’s multiple comparisons post hoc test). All data points are plotted in each graph, with black bars on top of data points that show the mean and standard deviation for each genotype. All images are inverted max projections unless specified. Scale bars = 5μm. CDS, Cell Division Site.

Next, we asked if ectopic Gef1 in *scd1Δ* cells and in LatA-treated cells leads to ectopic Cdc42 activation. We find that active Cdc42 appears depolarized in *scd1Δ* mutants during interphase. While CRIB-3xGFP remains restricted to the ends in *scd1+* cells, in *scd1Δ* mutants its localization appears as random patches all over the cortex (Figure 7Bi,ii). LatA treatment leads to ectopic CRIB-3xGFP (Mutavchiev et al., 2016) and this is abolished in *gef1Δ* cells (Figure 7Bvii,ix). In *gef1Δ* cells with LatA, CRIB-3xGFP remains at the cell ends, but at lower levels compared to untreated cells. This indicates that Scd1 is functionally epistatic to actin in the removal of Gef1 from the cell cortex. Together, these data demonstrate that Scd1 at the cells ends prevents ectopic Gef1 localization and Cdc42 activation.

Since we find that Scd1 regulates Gef1 localization at sites of polarized growth, we asked whether a similar relationship also occurs at the division site to facilitate Gef1 removal. To test this, we analyzed Gef1 localization in *scd1Δ* mutants. We observed persistent Gef1 localization in *scd1* mutants after ring constriction. In *scd1Δ* mutants, after completion of ring constriction and disassembly, Gef1 remains at the membrane that was adjacent to the ring (Figure 7C). Among *scd1Δ* cells that had completed constriction, 70% show persistent Gef1-mNG at the newly formed membrane barrier, as confirmed by the absence of Rlc1-tdTomato (Figure 7C,D). Similar Gef1-mNG localization was observed in only 20% of *scd1+* cells (Figure 7C,D, p<0.0001).

We find that Gef1-mNG removal from the division site is also dependent on actin. We analyzed Gef1 localization in cells with an actin cytoskeleton disrupted via Latrunculin A (LatA) treatment. In LatA-treated cells that were fully septated following completion of ring constriction, we observed persistent Gef1 localization at the division site. Gef1-mNG persists on both sides of the septum barrier in 40% of cells treated with LatA, but not in mock DMSO-treated cells (Figure 7E,F). Our data demonstrate that the actin cytoskeleton promotes Scd1 localization and that Scd1 promotes Gef1 removal from the division site. To confirm that actin promotes Gef1 removal via Scd1, we looked for a functional epistatic relationship between Scd1 and actin. We treated *scd1+* and *scd1Δ* cells expressing Gef1-mNG with LatA or DMSO. We find that in cells treated with DMSO, Gef1-mNG persists in 20% of septated *scd1+* cells and in 63% of septated *scd1Δ* cells. In cells treated with LatA, Gef1-mNG persists in 40% of septated *scd1+* cells and in 61% of septated *scd1Δ* cells (Figure 7F). The extent of Gef1 persistence in *scd1Δ* cells does not increase with the addition of LatA. This indicates that Scd1 is functionally epistatic to actin in the process of Gef1 removal (Figure 7F). Together, these data suggest that Scd1 removes Gef1 from the division site after ring disassembly.

To determine the significance of Scd1-mediated removal of Gef1 from the division site, we looked at the constitutively localized *gef1S112A* mutant. We confirmed that Gef1S112A persists at the division site after ring constriction. In control cells, Gef1-3xYFP is lost from the division site after the completion of ring constriction, indicated by the absence of Cdc15-tdTomato-marked actomyosin ring (Figure S8 A). However, Gef1S112A-3xYFP persists at the division site after ring constriction (Figure S8 A). Live cell imaging revealed that cell separation is delayed in *gef1S112A* mutants. Cell separation occurred 28 min after ring constriction in *gef1+* cells, but at 34 min in *gef1S112A* mutants (Figure S8 B, p=0.009). We have previously shown that prolonged Cdc42 activation at the division site impairs cell separation, as seen in other model systems (Atkins et al., 2013; Onishi et al., 2013; Wei et al., 2016). This would necessitate the removal of the GEFs from the division site. Our data indicate that Scd1-mediated Gef1 removal promotes cell separation.

## DISCUSSION

Polarity is essential for cell viability and development, as it regulates diverse processes such as motility, cell adhesion, secretion, determination of the division plane, and maintenance of cell fate. While studies in yeast continue to pioneer insights into the regulation of cell polarity, one aspect of S. *pombe* polarity remains elusive: how cells initiate growth at a second site during NETO (new end take-off). Since Cdc42 is the primary regulator of polarized growth, we examined its regulation during this process. We find that Gef1 enables the new end to overcome old end growth dominance to initiate bipolar growth. Our data indicate that Gef1 promotes the localization of the other GEF Scd1 to the new end. To uncover the mechanism by which this occurs, we examined the relationship between these two GEFs at the division site, where the localization of both these GEFs are easily monitored. We have recently shown that Gef1 and Scd1 localize sequentially to the division site to activate Cdc42 during cytokinesis (Wei et al., 2016). Here, we take advantage of the temporal difference between Gef1 and Scd1 localization at the division site to determine the significance of these two GEFs in Cdc42 regulation. We uncover a novel interplay between the Cdc42 GEFs that functions in both cytokinesis and polarized cell growth (Figure 8A). Given the conserved nature of Cdc42 and its regulators, we posit that this interplay between the GEFs is a common feature of Cdc42 regulation.

**Figure 8:**
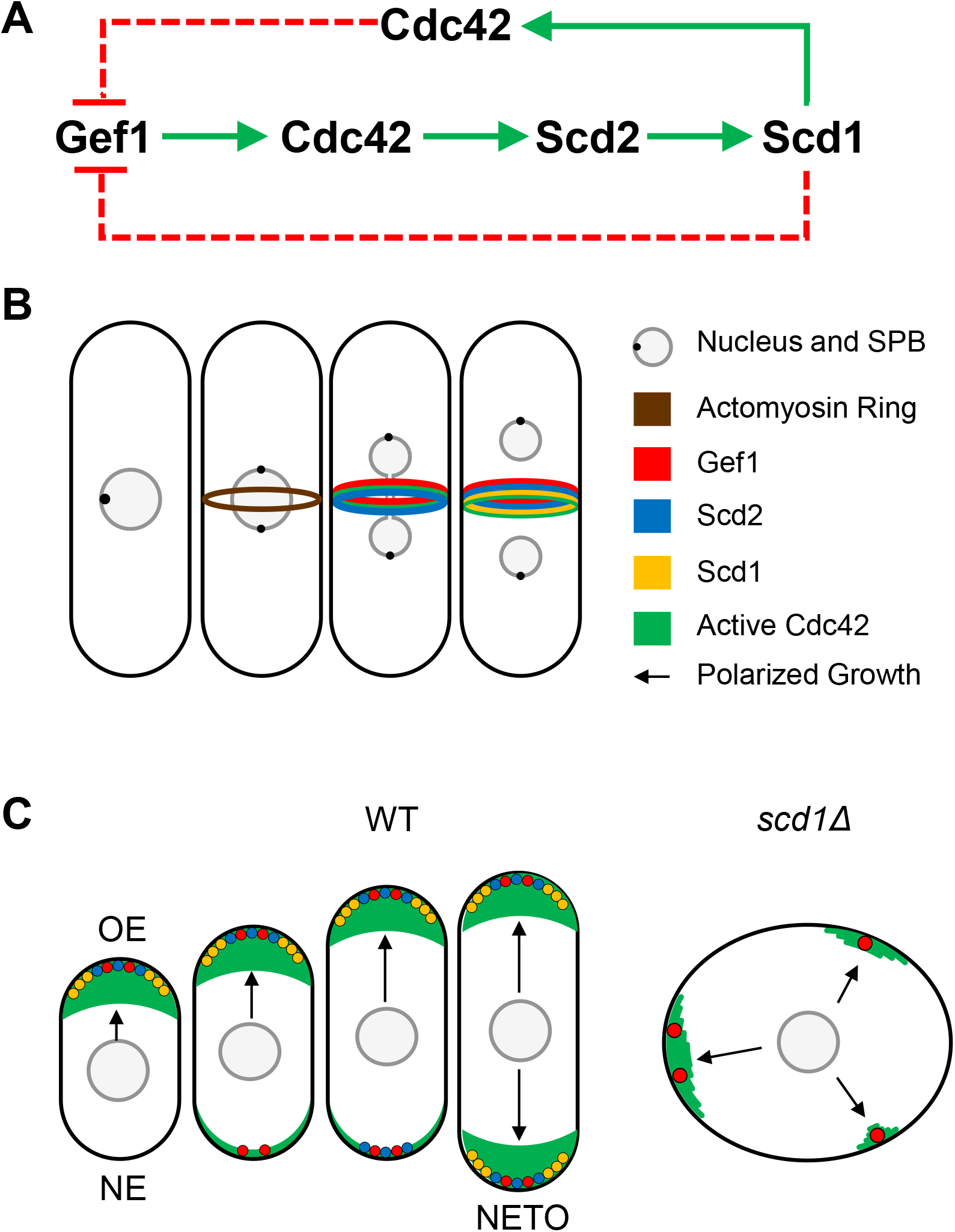
Model of the crosstalk between Gef1 and Scd1 that promotes polarized bipolar growth. **(A)** Diagram of the crosstalk pathway between Gef1 and Scd1. Solid arrows indicate an activating or promoting relationship in the direction of the arrow. Red terminating arrow indicates inactivation or removal of the protein at the arrows terminus. Dashed arrows indicate that the mechanism that regulates the proteins to which these arrows point is not yet resolved. **(B)** Schematic depicting the sequential localization of Gef1, Scd2, and Scd1 to the division site during cytokinesis. At the division site Gef1 localizes first and promotes Scd2 localization. Scd2 at the division site then recruits Scd1. **(C)** Schematic illustrating the crosstalk between Gef1 and Scd1 that promotes bipolar growth and regulates cell shape. In wild type (WT) cells, Gef1 activates Cdc42 which then recruits Scd2 to the new end, leading to Scd1 recruitment thus enabling NETO. In *scd1Δ* cells Gef1 localization is no longer restricted to the cell ends leading to ectopic Cdc42 activation and loss of polarity.

### Gef1-mediated Cdc42 activation promotes Scd1 recruitment via the scaffold Scd2

After division in fission yeast, the old end always initiates growth first. The two cell ends compete for active Cdc42, and initially the new end is incapable of overcoming dominance at the old end (Das et al., 2012). Gef1 has been shown to promote bipolar growth and cells lacking *gef1* are mostly monopolar (Coll et al., 2003; Das et al., 2012). Upon analysis of the growth pattern of *gef1Δ* mutants, we find that new ends frequently fail to overcome old end dominance, resulting in monopolar growth in these cells. Bipolar growth in *gef1Δ* mutants is typically observed in cells that do not contain a dominant old end. To determine how the new end overcomes old end growth dominance, we analyzed Cdc42 regulators in *gef1Δ* mutants. We find that Gef1 promotes bipolar localization of Scd1 to facilitate bipolar growth. Polarized growth requires Cdc42 activation, mediated through positive feedback (Das and Verde, 2013; Wu and Lew, 2013). In fission yeast, Scd1 is the positive-feedback-mediating GEF. Our data suggest that Gef1 activates Cdc42 to establish a Scd1-dependent positive feedback pathway at the new end to overcome old end dominance and establish bipolar growth (Figure 8C).

Analysis of Gef1-dependent Scd1 recruitment at the new end is complicated by the dynamic nature of their localization patterns, the presence of two competing cell ends, and the fact that these proteins do not localize to the cell ends in large quantities. Therefore, we investigated Gef1 and Scd1 recruitment at the division site. Gef1 and Scd1 localize to the division site in a sequential manner. Moreover, during cytokinesis, the division site exclusively activates Cdc42 and thus does not compete with any other site in the cell. We find that Scd1 localization is reduced at the division site in *gef1Δ* cells. Thus we posit that Gef1 localizes to the division site first, which enables recruitment of Scd1 (Figure 8B). Together with our observations at the cell ends, our data indicate that Gef1 promotes Scd1 localization via Cdc42 activation. Scd1 recruitment to the site of action is dependent on the scaffold Scd2. Scd1 and Scd2 together promote a positive feedback pathway for Cdc42 activation, where Scd2 binds active Cdc42. Once Scd1 activates Cdc42, it binds Scd2, which then recruits additional Scd1 to activate more Cdc42. For such a pathway to function, Cdc42 first needs to be activated for the recruitment of Scd2, and consequently Scd1. Our data indicate that Gef1-mediated Cdc42 activation provides the seed for Scd1-Scd2 recruitment. We find that Gef1 and Scd2 localize to the division site first, followed by Scd1. Similarly, ectopic Gef1 localization as observed in LatA-treated cells shows complete colocalization with Scd2, suggesting that Gef1-mediated Cdc42 activation promotes Scd2 recruitment. This enables Scd2 recruitment to nascent sites that have no prior history of Cdc42 activation and helps Scd1 recruitment to those sites.

### Scd1 prevents ectopic Gef1 localization

In fission yeast, Scd1 is the primary GEF that promotes polarized growth (Chang et al., 1994). We find that cells lacking *scd1* are depolarized due to ectopic Cdc42 activation, as a result of mislocalized Gef1. In the presence of *scd1*, Gef1 shows sparse cortical localization and is restricted to the cell ends. We find that cell ends that do not localize Scd1 either due to LatA treatment or due to *scd1Δ* display ectopic and enhanced cortical localization if Gef1. Thus we posit that Scd1 prevents ectopic Gef1 localization. A recent report shows that ectopic Cdc42 activation in LatA-treated cells depends on the stress-activated MAP kinase Sty1 (Mutavchiev et al., 2016). Fission yeast cells treated with LatA did not display ectopic Cdc42 activation in the absence of *sty1.* It is possible that in the absence of actin, reduced Scd1 at the cell cortex elicits a stress response, leading to Sty1 activation that results in the mislocalization of Gef1. Further analysis is necessary to test this hypothesis.

Similar to our observation at the cell cortex, we find that Gef1 persists at the division site in cells lacking Scd1 or in cells treated with LatA. Normally, Gef1 localization to the division site is lost after ring constriction (Wei et al., 2016). Here, we show that Scd1 promotes the clearance of Gef1 from the division site after ring disassembly (Figure 8A). Mis-regulation of Cdc42 has been reported to result in cytokinesis failure in many organisms. Specifically, failure to inactivate Cdc42 leads to failed cell abscission in budding yeast and HeLa cells, and prevents cellularization in *Drosophila* embryos (Atkins et al., 2013; Crawford et al., 1998; Dutartre et al., 1996; Onishi et al., 2013). The mechanism by which Cdc42 is inactivated prior to cell abscission is unclear. Our data suggest that Scd1 ensures that Gef1 does not persist at the division site in the final stages of cytokinesis, preventing inappropriate Cdc42 activation. Together, our data demonstrate an elegant regulatory pattern in which Gef1-mediated Scd1 recruitment to the division site promotes septum formation, and Scd1-mediated Gef1 removal promotes cell separation.

### Multiple GEFs combinatorially regulate Cdc42 during complex processes

Polarized cell growth requires symmetry breaking, and several models have indicated a need for Cdc42 positive feedback loops in this process (Bendezu et al., 2015; Irazoqui et al., 2003; Kozubowski et al., 2008; Slaughter et al., 2009a; Slaughter et al., 2009b; Wedlich-Soldner et al., 2004). Elegant experiments in budding yeast demonstrate that local activation of Cdc42 establishes positive feedback through the recruitment of additional GEFs to amplify the conversion of Cdc42-GDP to Cdc42-GTP (Butty et al., 2002; Kozubowski et al., 2008). A caveat of positive feedback is that the site that first activates Cdc42 can act as a sink that traps the GEFs, thereby preventing Cdc42 activation at other sites. Such a trap can be undone via negative feedback regulation of Cdc42 that results in an oscillatory pattern at the cell ends (Das et al., 2012; Howell et al., 2012). Negative feedback has reported in S. *pombe* and S. *cerevisiae* through the Pak1 kinase activity that antagonizes either the Cdc42 scaffold or the GEF (Das et al., 2012; Gulli et al., 2000; Kuo et al., 2014; Rapali et al., 2017). Indeed, *pak1* mutants fail to activate bipolar growth (Das et al., 2012; Verde et al., 1998). Our data show that in addition to the Pak1 kinase, Gef1 also contributes to initiation of bipolar growth.

The Cdc42 oscillatory pattern can be explained by the presence of positive feedback, time-delayed negative feedback, and competition between the two ends for active Cdc42. Since Scd1 is the Cdc42 GEF that establishes polarized growth, we posit that Scd1 activates Cdc42 through positive feedback at the dominant old end. Dominance at the old end ensures that Scd1 localization is mainly restricted to this end at the expense of the new end. A previous model suggests that as the cell reaches a certain size, the GEFs reach a threshold level that allows the new end to overcome old end dominance to initiate growth and promote bipolarity (Das et al., 2012). Threshold GEF levels alone cannot explain our findings since *gef1S112A* cells display bipolar growth at a smaller cell size (Das et al., 2015), while G1-arrested *cdc10-129* mutants grow to longer cell lengths but remain monopolar. Our data suggest that the regulatory crosstalk between the Cdc42 GEFs may provide an advantage to the cell and enables the new end to overcome old end dominance. Gef1 activates Cdc42 at the new end that then recruits scd2 to finally recruit Scd1. Thus Gef1 triggers a positive feedback at the division site via a feed-forward pathway (Figure 8A,C). Given that Gef1 promotes Scd1-mediated polarized growth at the new end, it is conceivable that Gef1 itself is tightly regulated to prevent random Cdc42 activation. Indeed, Gef1 shows sparse localization to the cell ends and is mainly cytoplasmic (Das et al., 2015). The NDR kinase Orb6 prevents ectopic Gef1 localization via 14-3-3-mediated sequestration to the cytoplasm (Das et al., 2015; Das et al., 2009). Here we show that while Gef1 promotes Scd1 recruitment to a nascent site, Scd1 itself restricts Gef1 localization to the cell ends to precisely activate Cdc42 (Figure 8C). Together, our findings describe an elegant system in which the two Cdc42 GEFs regulate each other to ensure proper cell polarization.

### Significance of GEF coordination in other systems

In budding yeast, CDC24 is required for polarization during bud emergence and is essential for viability (Sloat et al., 1981; Sloat and Pringle, 1978), unlike Scd1 in fission yeast. Budding yeast also has a second GEF Bud3, which establishes a proper bud site (Kang et al., 2014). During G1 in budding yeast, bud emergence occurs via biphasic Cdc42 activation by the two GEFs: Bud3 helps select the bud site (Kang et al., 2014), and Cdc24 allows polarization (Sloat et al., 1981; Sloat and Pringle, 1978). This is analogous to new end growth in fission yeast, which requires Gef1-dependent recruitment of Scd1 for robust Cdc42 activation. It would be interesting to test whether crosstalk also exists between Bud3 and Cdc24.

The Rho family of GTPases includes Rho, Rac, and Cdc42. In certain mammalian cells, Cdc42 and Rac1 appear to activate cell growth in a biphasic manner (de Beco et al., 2018; Yang et al., 2016). For example, during motility, the GTPases, Rho, Rac, and Cdc42, regulate the actin cytoskeleton (Heasman and Ridley, 2008; Machacek et al., 2009). During cell migration, these GTPases form bands or ‘zones’ in the leading and trailing regions of the cell (Ridley, 2015).

Their spatial separation is mediated by the organization of their GEFs and GAPs, as well as by regulatory signaling between these GTPases (Guilluy et al., 2011). Cdc42 and Rho are mutually antagonistic, explaining how such zones of GTPase activity can be established and maintained (Guilluy et al., 2011; Kutys and Yamada, 2014; Warner and Longmore, 2009). Similarly, Cdc42 can refine Rac activity (Guilluy et al., 2011). Cdc42 and Rac are activated by similar pathways and share the same effectors. Several recent experiments demonstrate that during cell migration, reorganization of the actin cytoskeleton occurs in a biphasic manner, in which Cdc42 activation at new sites sets the direction, while robust Rac activation determines the speed (de Beco et al., 2018; Yang et al., 2016). Unlike most eukaryotes, the genome of S. *pombe* does not contain a Rac GTPase. We speculate that the two Cdc42 GEFs of *S. pombe* allow it to fulfill the roles of both Cdc42 and Rac. Gef1 sets the direction of growth by establishing growth at a new site, while Scd1 promotes efficient growth through robust Cdc42 activation at the growth sites. In conclusion, we propose that the crosstalk between the Cdc42 GEFs themselves is an intrinsic property of small GTPases and is necessary for fine-tuning their activity.

## MATERIALS AND METHODS

### Strains and cell culture

The S. *pombe* strains used in this study are listed in Supplemental Table S1. All strains are isogenic to the original strain PN567. Cells were cultured in yeast extract (YE) medium and grown exponentially at 25°C, unless specified otherwise. Standard techniques were used for genetic manipulation and analysis (Moreno et al., 1991). Cells were grown exponentially for at least 3 rounds of eight generations each before imaging.

### Microscopy

Cells were imaged at room temperature (23–25°C) with an Olympus IX83 microscope equipped with a VTHawk two-dimensional array laser scanning confocal microscopy system (Visitech International, Sunderland, UK), electron-multiplying charge-coupled device digital camera (Hamamatsu, Hamamatsu City, Japan), and 100×/numerical aperture 1.49 UAPO lens (Olympus, Tokyo, Japan). Images were acquired with MetaMorph (Molecular Devices, Sunnyvale, CA) and analyzed by ImageJ (National Institutes of Health, Bethesda, MD).

### Analysis of growth pattern

The growth pattern of *gef1+* and *gef1Δ* cells was observed by live imaging of cells through multiple generations. Cells were placed in 3.5-mm glass-bottom culture dishes (MatTek, Ashland, MA) and overlaid with YE medium plus 1% agar, and 100μM ascorbic acid to minimize photo-toxicity to the cell. A bright-field image was acquired every minute for 12 hours. Birth scars were used to distinguish between, as well as to measure, old end and new end growth.

### Construction of fluorescently tagged Gef1 fusion proteins

The forward primer 5’-GGATCCGTGTTTACCAAAGTTATGTAAGAC -3’ with a 5’ BamHI site and the reverse primer 5’-CCCGGGAACCCTCGCAGCTAAAGA -3’ with a 5’ Xmal site were used to amplify a 3kb DNA fragment containing *gef1*, the 5’ UTR, and the endogenous promoter. The fragment was then digested with BamHI and XmaI and ligated into the BamHI-XmaI site of pKS392 pFA6-tdTomato-kanMX and pKG6507 pFA6-mNeonGreen-kanMX. Constructs were linearized by digestion with XbaI and transformed into the *gef1* locus in *gef1Δ* cells.

### Expression of constitutively active Cdc42

*pjk148-nmt41x-leu1^+^* or *pjk148-nmt41x:cdc42G12V-leu1^+^* were linearized with NdeI and integrated into the *leu1-32* locus in *gef1+* and *gef1Δ* cells expressing either CRIB-3xGFP and Scd1-tdTomato, or Scd1-tdTomato and Rlc1-GFP. The empty vector *pjk148-nmt41x-leu1^+^*was used as control. Cells were grown in EMM with 0.05uM thiamine to promote minimal expression of *cdc42G12V.*

### Latrunculin A treatment

Cells in YE were incubated at room temperature with 10μM or 100 μM Latrunculin A (Millipore-Sigma) dissolved in dimethyl sulfoxide (DMSO) for 40 min prior to imaging. Control cells were treated with 1% DMSO and incubated for 40 min.

### Analysis of fluorescent intensity

Mutants expressing fluorescent proteins were grown to OD 0.5 and imaged on slides. Cells in slides were imaged for no more than 3 minutes to prevent any stress response as previously described (Das et al., 2015). Depending on the mutant and the fluorophore, 16-28 Z-planes were collected at a z-interval of 0.4μm for either or both the 488nm and 561nm channels. The respective controls were grown and imaged in an identical manner. ImageJ was used to generate sum projections from the z-series, and to measure the fluorescence intensity of a selected region (actomyosin ring, or growth cap at cell tip). The background fluorescence in a cell-free region of the image was subtracted to generate the normalized intensity. Mean normalized intensity was calculated for each image from all (n>5) measurable cells within each field. A Student’s two-tailed t-test, assuming unequal variance, was used to determine significance through comparison of each strain’s mean normalized intensities.

## Supporting information

Supplemental Figures and strains list

## Acknowledgements

We thank J. Bembenek and T. Burch-Smith for critical review of our manuscript; K. Gould for supplying plasmids; and M. Balasubramanian, and S. Martin for providing strains.

