## Supplemental Figures and strains list for "A novel interplay between GEFs orchestrates Cdc42 activity during cell polarity and cytokinesis"

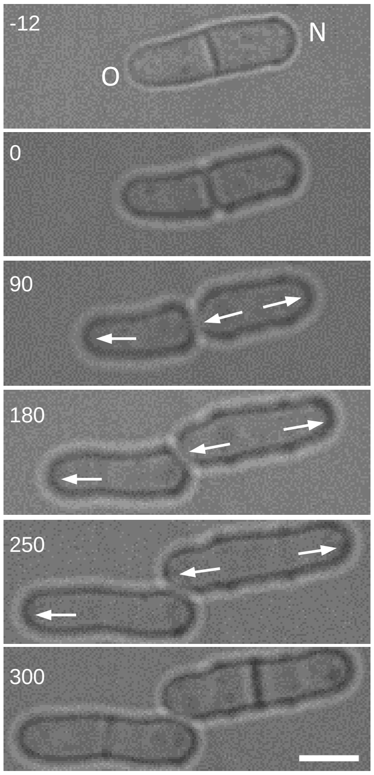


**Supplemental Figure 1: Growth pattern in the progeny of a monopolar *gef1Δ* cell.** Image at -12 mins shows a monopolar *gef1Δ* cell that in the previous generation grew only from the old end (O) while the new end (N) did not have any prior history of growth. Time stamps are minutes elapsed since completion of division. Arrows indicate direction of growth at cell ends. Scale bar=5µm.

Supplemental Figure 1

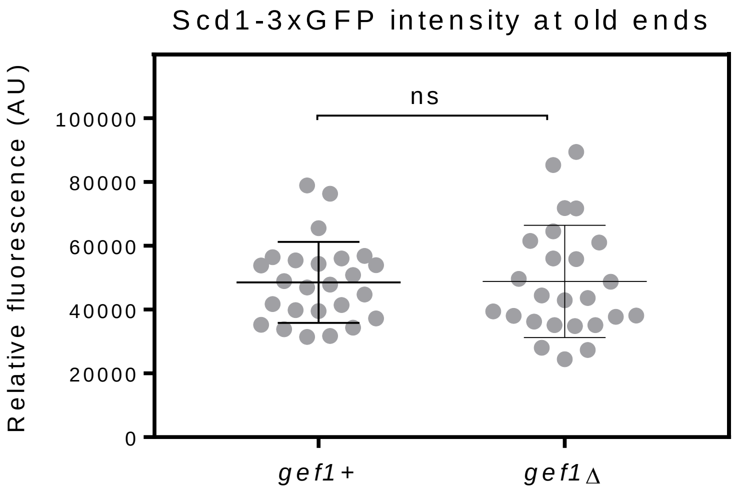


**Supplemental Figure 2: Scd1 localization to the old end is not impaired in *gef1Δ*.** Quantification of Scd1-3xGFP localization to old ends in *gef1+* and *gef1Δ cells*. n.s, not significant.

Supplemental Figure 2

Scd1-3xGFP

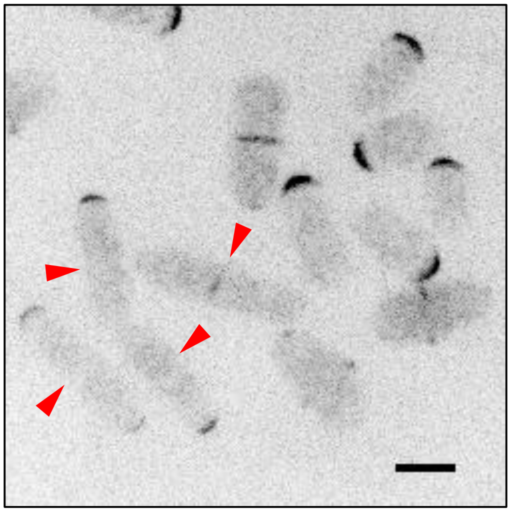

**Supplemental Figure 3: The PAK kinase antagonizes Scd1 accumulation to limit Cdc42 activity.** Scd1-3xGFP accumulation in *pak1+* and nmt1:*pak1 switch-off* mutant cells. Cells were grown to an OD of 0.5 in minimal media + thiamine and mixed prior to imaging. Red arrow heads indicate *pak1+* cells. Images are inverted max projections. Scale bar=5µm.

Supplemental Figure 3

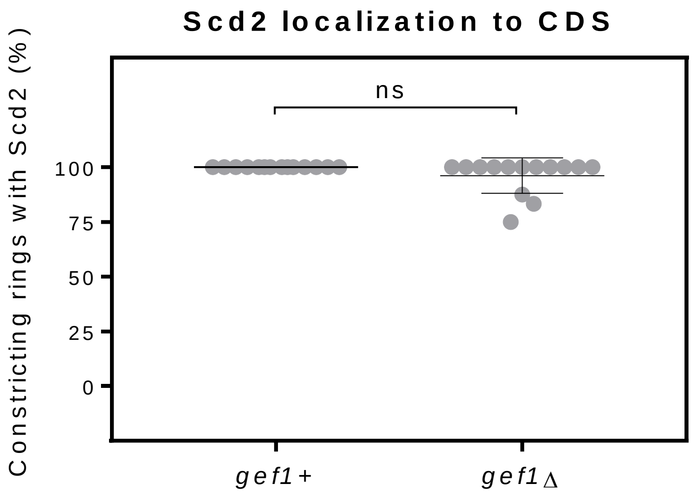


**Supplemental Figure 4: Scd2 localization to the division site is delayed until the onset of ring constriction in *gef1Δ*.** Quantification of Scd2-GFP localization to constricting rings in *gef1+* and *gef1Δ*. (ns, not significant). Cell division site, CDS.

Supplemental Figure 4

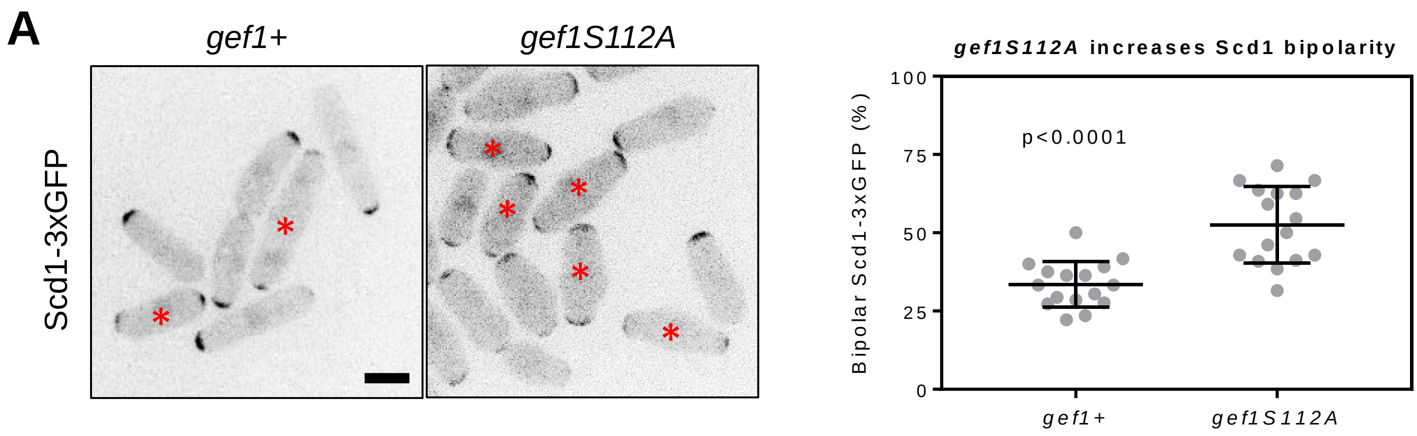


**Supplemental Figure 5: Gef1 promotes the transition to bipolar growth. (A)** Localization of Scd1-3xGFP to the cell poles in *gef1+* and *gef1S112A* cells. Asterisks indicate cells with bipolar Scd1-3xGFP localization. **(B)** Quantification of the percent of cells that exhibit bipolar Scd1-3xGFP localization at cell ends in the indicated genotypes. Scale bar=5µm.

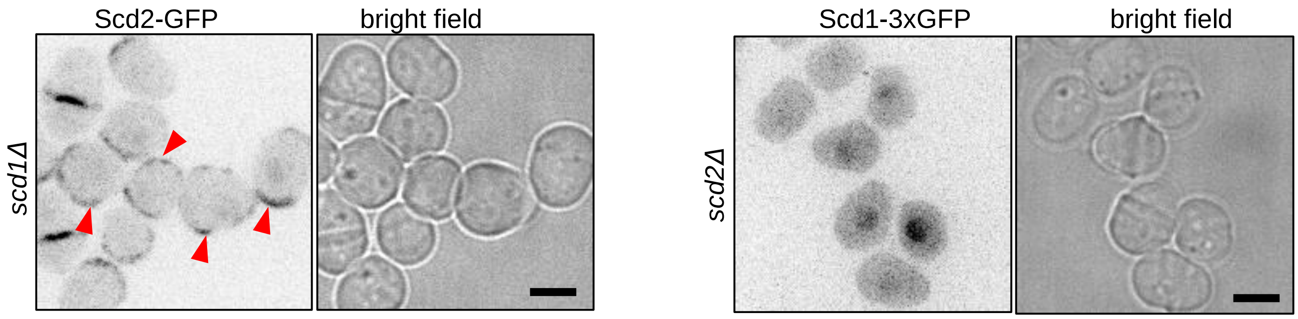


**Supplemental Figure 6: Scd1 requires *scd2* to localize to the CDS and the cortex, but Scd2 localization is unimpaired in the absence of *scd1*.** Scd2-GFP and Scd1-3xGFP localization to the cortex in *scd1Δ* and *scd2Δ* cells, respectively. Red arrowheads indicate cells with Scd2-GFP localized tot the cell cortex. All images are inverted max projections with the exception of bright. Scale bars = 5µm. Cell Division Site, CDS.

Supplemental Figure 6

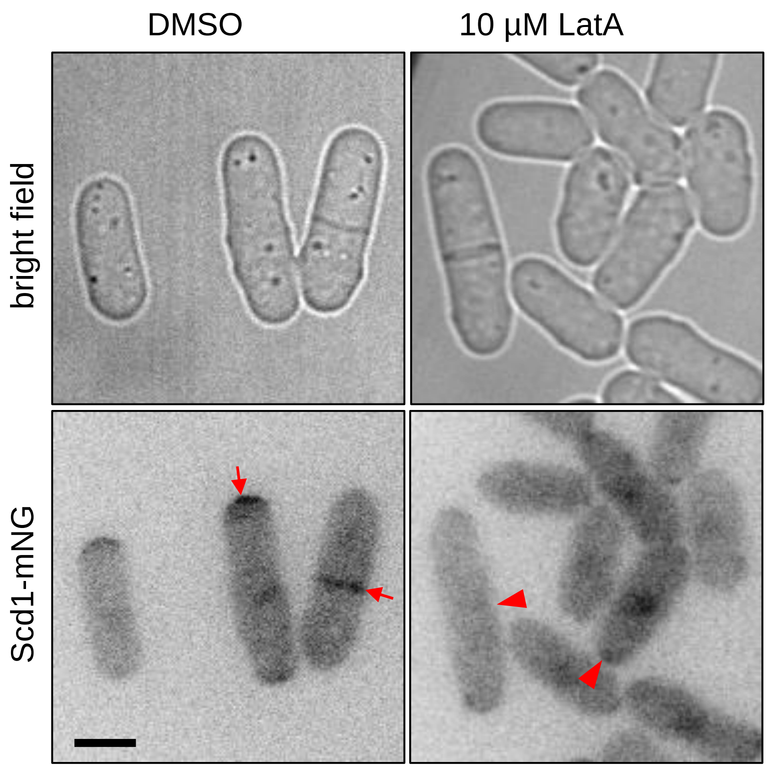


**Supplemental Figure 7: Scd1 requires actin for its localization to the CDS and the cortex.** Scd1-mNG localization to the division site and cell tips in cells treated with 10 µM LatA or DMSO. Red arrows indicate cells with Scd1-mNG localized to the CDS or cell tip. Red arrowheads indicate cells where Scd1-mNG fails to localize to the division site or to the cell tips. All images are inverted max projections with the exception of bright field. Scale bars = 5µm. Cell Division Site, CDS.

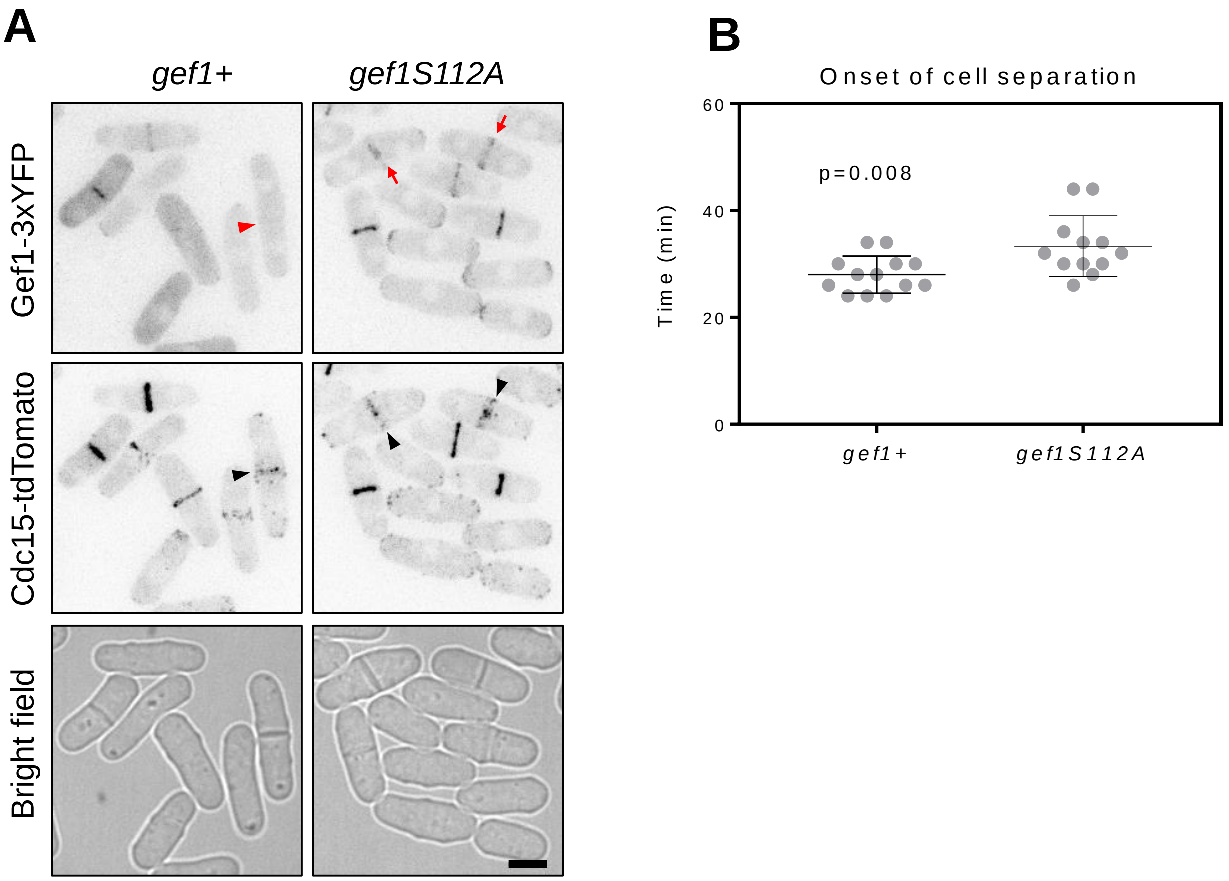


**Supplemental Figure 8: Gef1 removal form the division site promotes cell separation. (A)** Gef1-3YFP and Gef1S112A-3YFP localization in cells expressing the ring marker Cdc15-tdTomato. Red arrow head indicates the absence of Gef1 from the division site after ring constriction. Red arrows indicate the presence of Gef1 at the division site after ring constriction. Black arrow heads mark the division site of cells that have completed ring constriction. **(B)** Quantification of onset of cell separation after completion of ring constriction in *gef1+* and *gef1S112A* cells. Scale bar=5µm.

Supplemental Figure 8

| **Table S1. Strains list.** | | |
|---|---|---|
| Strain | Genotype | Source |
| PN567 | h+ ade6-704 leu1-32 ura4-d18 | Paul Nurse |
| PN1191 | cdc10-129 ade6-704 leu1-32 | Paul Nurse |
| JX125 | h90 Δscd1::ura4+ ade6 leu1-32 ura4-d18 h210 | (Hirota et al., 2003) |
| FV1218 | Gef1-3xYFP:kanMX ade6-704 leu1-32 ura4-d18 | (Das et al., 2009) |
| YSM947 | Scd1-3xGFP:kanMX ade6-m216 leu1-32 ura4-d18 | (Bendezu and Martin, 2013) |
| MBY3451 | P3 nmt1:3HA-Shk1:KANr leu1-32 ura4-d18 | (Loo and Balasubramanian, 2008) |
| YMD1172 | Scd1-tdTomato:KANr ade6-704 leu1-32 ura4-d18 | This study |
| YMD1256 | Δgef1::ura4+ pjk148-nmt41x:cdc42G12V-leu1+ CRIB-3xGFP:ura4+ Scd1-tdTomato:KanMX ade6 leu1-32 ura4-d18 | This study |
| YMD1271 | pjk148-nmt41x-leu1+ CRIB-3xGFP:ura4+ Scd1-tdTomato:KanMX ade6 leu1-32 ura4-d18 | This study |
| YMD1273 | Δgef1::ura4+ pjk148-nmt41x-leu1+ CRIB-3xGFP:ura4+ Scd1-tdTomato:KanMX ade6 leu1-32 ura4-d18 | This study |
| YMD1232 | pjk148-nmt41x:cdc42G12V-leu1+ CRIB-3xGFP:ura4+ Scd1-tdTomato:KanMX ade6 leu1-32 ura4-d18 | This study |
| YMD317 | CRIB-3xGFP:ura4+ Rlc1-tdTomato:NATr Sad1-mCherry:kanMX ade6-M21X leu1-32 ura4-D18 | (Wei et al., 2016) |
| YMD432 | Δgef1::ura4+ pjk148-nmt41x:cdc42G12V-leu1+ CRIB-3xGFP:ura4+ ade6 leu1-32 ura4-d18 | This study |
| YMD488 | Δgef1::ura4+ CRIB-3xGFP:ura4+ Rlc1-tdTomato:NATr Sad1-mCherry:kanMX ade6 leu1-32 ura4-d18 | (Wei et al., 2016) |
| YMD530 | h90 Δscd1::ura4+ CRIB-3xGFP:ura4+ Rlc1-tdTomato:NATr Sad1-mCherry:kanMX ade6-M21X leu1-32 ura4-d18 | (Wei et al., 2016) |
| YMD602 | pjk148-nmt41x:cdc42G12V-leu1+ CRIB-3xGFP:ura4+ ade6 leu1-32 ura4-d18 | This study |
| YMD1043 | Gef1-tdTomato:KANr Scd2-GFP:KANr ade6 leu1-32 ura4-d18 | This study |
| YMD1204 | Scd1-tdTomato:KANr Scd2-GFP:KANr ade6 leu1-32 ura4-d18 | This study |

| | | |
|---|---|---|
| YMD1049 | Scd1-3xGFP:KANr Gef1-tdTomato:KANr ade6 leu1-32 ura4-d18 | This study |
| YMD761 | Δgef1::ura4+ Scd1-3xGFP:kanMX Rlc1-tdTomato:NATr Sad1-mCherry:kanMX ade6-m216 leu1-32 ura4-d18 | This study |
| YMD773 | Scd1-3xGFP:kanMX Rlc1-tdTomato:NATr Sad1-mCherry:kanMX ade6-m216 leu1-32 ura4-d18 | This study |
| YMD795 | nmt1:3HA-Shk1 scd2-GFP:kanMX | This study |
| YMD840 | Δgef1::ura4+ Scd2-GFP:kanMX Rlc1-tdTomato:NATr Sad1-mCherry:kanMX ade6 leu1-32 ura4-d18 | This study |
| YMD842 | Scd2-GFP:kanMX Rlc1-tdTomato:NATr Sad1-mCherry:kanMX kanMX ade6 leu1-32 ura4-d18 | This study |
| YMD910 | Gef1-NeonGreen:kanMX leu1-32 ura4-d18 | This study |
| YMD926 | Gef1-NeonGreen:kanMX Rlc1-tdTomato:NATr Sad1-mCherry:kanMX ade6 leu1-32 ura4-d18 | This study |
| YMD936 | Δgef1::ura4+ pjk148-nmt41x:cdc42G12V-leu1+ Scd1-3xGFP:kanMX ade6-m216 leu1-32 ura4-d18 | This study |
| YMD994 | pjk148-nmt41x:cdc42G12V-leu1+ Scd1-3xGFP:kanMX ade6-m216 leu1-32 ura4-d18 | This study |
| YMD996 | h90 Δscd1::ura4+ Scd2-GFP:kanMX Rlc1-tdTomato:NATr ade6 leu1-32 ura4-d18 | This study |
| YMD998 | pjk148-nmt41x-leu1+ CRIB-3xGFP:ura4+ ade6 leu1-32 ura4-d18 | This study |
| YMD1000 | Δgef1::ura4+ pjk148-nmt41x-leu1+ CRIB-3xGFP:ura4+ ade6 leu1-32 ura4-d18 | This study |
| YMD1002 | Δgef1::ura4+ pjk148-nmt41x-leu1+ Scd1-3xGFP:kanMX ade6-m216 leu1-32 ura4-d18 | This study |
| YMD1004 | pjk148-nmt41x-leu1+ Scd1-3xGFP:kanMX ade6-m216 leu1-32 ura4-d18 | This study |
| YMD1030 | h90 Δscd1::ura4+ Gef1-NeonGreen:kanMX Rlc1-tdTomato:NATr Sad1-mCherry:kanMX ade6 leu1-32 ura4-d18 | This study |
| YMD1067 | h90 Δscd2::ura4+ Gef1-NeonGreen:kanMX Sad1-mCherry:kanMX ade6 leu1-32 ura4-d18 h210 | This study |
| YMD1069 | h90 Δscd2::ura4+ Scd1-3xGFP:kanMX Rlc1-tdTomato:NATr ade6-m216 leu1-32 ura4-d18 | This study |
| YMD1088 | gef1s112a:kanMX Scd1-3xGFP:kanMX ade6-m216 leu1-32 ura4-d18 | This study |
| YMD965 | Δgef1::ura4+ gef1S112A-3xYFP:kanMX Cdc15-tdTomato:$Nat^r$ ade6 leu1-32 ura4-d18 | This study |

| | | |
|---|---|---|
| YMD208 | Cdc15-tdTomato:NAT$^r$ Gef1-3xYFP:kanMX ade6 | (Wei et al., 2016) |
